# Defining a small-molecule stimulator of the human Hsp70-disaggregase system with selectivity for DnaJB proteins

**DOI:** 10.1101/2024.01.11.575109

**Authors:** Edward Chuang, Ryan R. Cupo, Jeffrey R. Bevan, Mikhaila L. Rice, Shuli Mao, Erica L. Gorenberg, Korrie L. Mack, Donna M. Huryn, Peter Wipf, Jeffrey L. Brodsky, James Shorter

## Abstract

Hsp70, Hsp40, and Hsp110 form a human protein-disaggregase system that solubilizes and reactivates proteins trapped in aggregated states. However, this system fails to maintain proteostasis in fatal neurodegenerative diseases. Here, we potentiate the human Hsp70-disaggregase system pharmacologically. By scouring a collection of dihydropyrimidines, we disambiguate a small molecule that specifically stimulates the Hsp70-disaggregase system against disordered aggregates and α-synuclein fibrils. The newly identified lead compound stimulates the disaggregase activity of multiple active human Hsp70, Hsp40, Hsp110 chaperone sets, with selectivity for combinations that include DnaJB1 or DnaJB4 as the Hsp40. We find that the relative stoichiometry of Hsp70, Hsp40, and Hsp110 dictates disaggregase activity. Remarkably, our lead compound shifts the composition of active chaperone stoichiometries by preferentially activating combinations with lower DnaJB1 concentrations. Our findings unveil a small molecule that stimulates the Hsp70-disaggregase system, even at suboptimal chaperone stoichiometries, which could be developed for the treatment of neurodegenerative diseases.

**Graphical Abstract:** 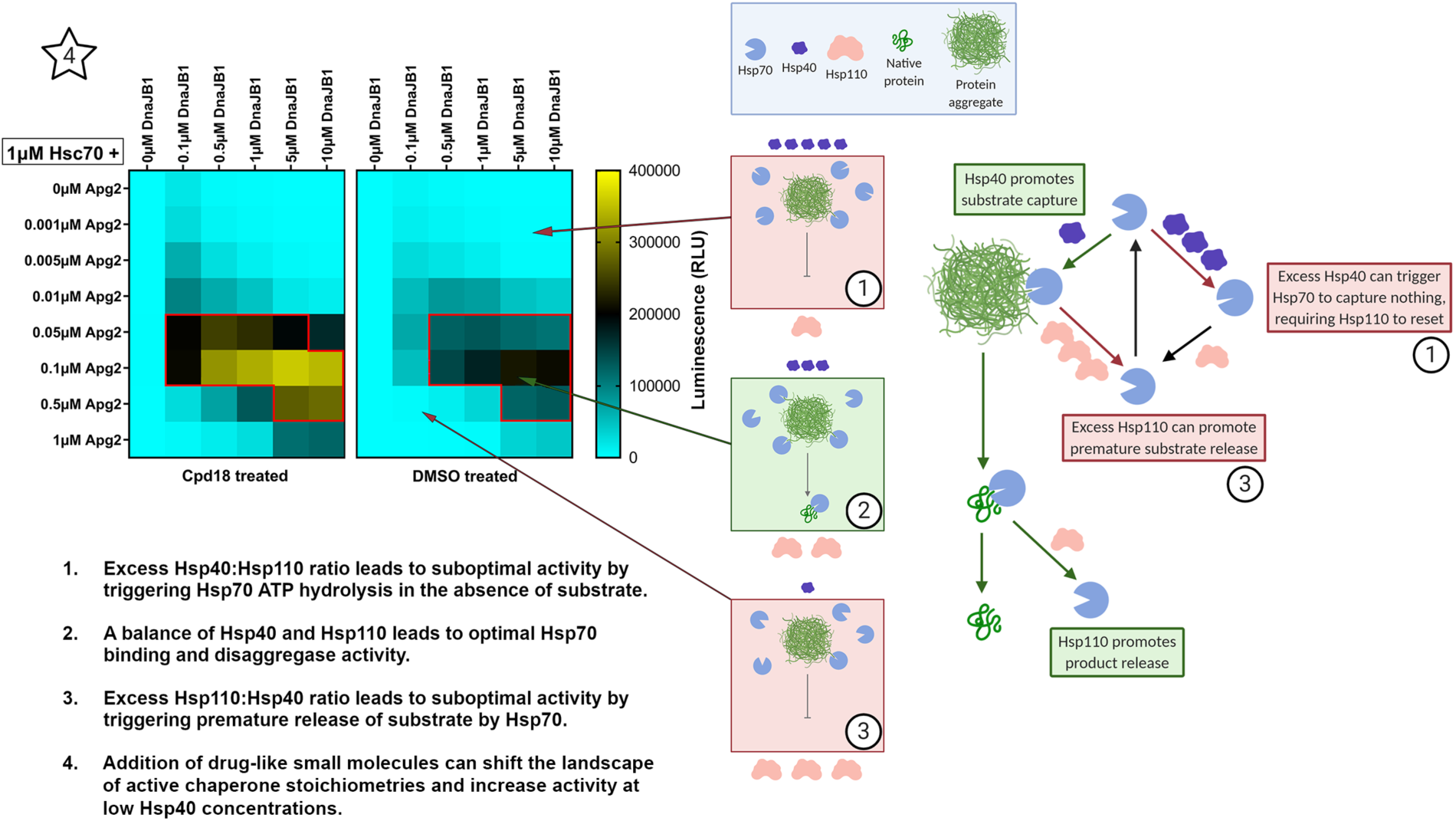

## Introduction

Proteins must fold properly to perform a myriad of functions.^1,2^ During stress, proteins may become misfolded and aggregate through aberrant intra-and intermolecular interactions or remain folded but become trapped in phase-separated states.^3–5^ Some proteins are particularly prone to aggregation and accumulate in the brains of patients with neurodegenerative diseases.^6^ For example, in degenerating neurons of Parkinson’s disease (PD) patients, α-synuclein (αSyn) forms insoluble inclusions in the cytoplasm called Lewy bodies.^6,7^ Similar hallmarks of protein aggregation are observed in other neurodegenerative diseases such as Alzheimer’s disease, amyotrophic lateral sclerosis (ALS), and frontotemporal dementia (FTD).^6,7^

Protein aggregates, amyloids, and their oligomeric precursors can be toxic by conferring gain-of-function and loss-of-function phenotypes.^8^ Devising ways to remove these toxic conformers and restore proteins back to native form and function could present an avenue to treat neurodegenerative diseases.^6,8^ One possible strategy is to leverage the sophisticated protein disaggregases that cells have evolved to disaggregate and reactivate proteins trapped in aberrant states.^8^ Yet these systems fail in neurodegenerative disease. Thus, stimulating the activity of endogenous protein disaggregases may provide a mechanism to counter deleterious protein-misfolding events that underlie neurodegenerative disease.^9,10^

The 70 and 40 kDa heat shock proteins (Hsp70 and Hsp40) form one of the predominant molecular chaperone systems that unfold and refold misfolded proteins.^11^ However, aggregated proteins contain stable intermolecular interactions that can be difficult to break.^12^ Hsp110, a member of the Hsp70 super-family, collaborates with Hsp70 and Hsp40 to enable the disassembly of protein aggregates and restoration of native protein function.^13–17^ Specifically, the human Hsp70-disaggregase system, comprising Hsp70, Hsp40, and Hsp110 family members, can disassemble disordered aggregates such as urea-denatured luciferase and heat-denatured GFP *in vitro*^13,16,18^ as well as ordered Sup35, αSyn, Huntingtin(Htt)-polyQ, and tau amyloid fibrils.^19–28^ Hsp70 chaperone activity requires controlled binding and release of protein substrates.^29,30^ In the open ATP-bound conformation, polypeptides can bind to the substrate-binding domain (SBD) of Hsp70 with a low affinity and a high exchange rate.^30–32^ In the closed ADP-bound conformation, substrate is trapped in the SBD with a high affinity and a low exchange rate.^30,33^ In turn, the ATP cycle of Hsp70 is regulated by Hsp40 and Hsp110.^14,29,30^ Hsp40 binds to substrate and recruits it to Hsp70.^14,29,30^ Concomitant binding of Hsp40 and substrate to Hsp70 promotes Hsp70 ATP hydrolysis resulting in substrate capture.^14,29,30^ Hsp110 is a nucleotide-exchange factor (NEF) for Hsp70 and promotes exchange of ADP for ATP, thus reverting Hsp70 back to the open state.^6,13,14,34–39^ Thus, through coordination with Hsp40 and Hsp110, Hsp70 can bind and extract a polypeptide from an aggregate, and then release the polypeptide, allowing it to refold into its native conformation.^6,14,29,40,41^

The human genome contains 12 Hsp70-encoding genes, 55 Hsp40-encoding genes, and four Hsp110-encoding genes, giving rise to thousands of potential three-component combinations of the human Hsp70-disaggregase system.^14,42^ Lower-order organisms, such as bacteria or yeast, encode significantly fewer members of these proteins in their genomes. In *E. coli* there are three Hsp70 genes, six Hsp40 genes, and one Hsp70 NEF gene, which is not an Hsp110 homolog. In *S. cerevisiae* there are 11 Hsp70 genes, 22 Hsp40 genes, and two Hsp110 genes.^14^ It is hypothesized that expanded cohorts of Hsp70, Hsp40, and Hsp110/NEF genes results in a non-linear increase in the number of unique chaperone combinations, which may enable a limited number of chaperones to survey significantly larger proteomes with greater specificity.^14,43^ Furthermore, the Hsp40 protein family can be categorized into class A, class B, and class C J-domain proteins.^44^ It has been reported that the class A and class B Hsp40 proteins can synergize to yield greater luciferase disaggregase activity for chaperone sets composed of Hsp70, Hsp110, and two Hsp40 members.^15^ Thus, the human Hsp70-disaggregase system can be formed by three or four component sets, greatly increasing the combinatorial space of the chaperone network.

One select combination, Hsc70, DnaJB1, and Apg2 can disassemble αSyn amyloid fibrils more effectively than other combinations, suggesting that different combinations of the Hsp70 system can have drastically different efficacy against the same substrate.^20^ This same combination also disaggregates Htt-PolyQ fibrils and tau fibrils.^21,22^ These findings suggest that an endogenous disaggregase machinery in human cells can robustly disaggregate protein aggregates *in vitro*, and yet fails in the brains of patients with neurodegenerative diseases. Thus, is the Hsp70-disaggregase system a therapeutic target? That is, could we find small-molecule drugs that stimulate the Hsp70-disaggregase system to restore proteostasis and counter protein misfolding in neurodegenerative disease?^9,45^

One small molecule, 115-7c (also known as MAL1-271), is a dihydropyrimidine that stimulates the ATPase and protein-folding activity of prokaryotic Hsp70 (DnaK).^46–49^ 115-7c binds at the interface between the nucleotide-binding domain (NBD) of DnaK and the J-domain of DnaJ (prokaryotic Hsp40) and is proposed to stabilize the interaction between DnaK and DnaJ, thus promoting ATP hydrolysis and substrate capture.^49^ The proposed mechanism for the human Hsp70 system shares the same ATP hydrolysis and substrate capture step, which is promoted by 115-7c.^6,14^ Thus, we might be able to target the human Hsp70-disaggregase system with 115-7c or related scaffolds to bolster the disaggregase machinery within patient neurons to combat aberrant protein aggregation.^9,45^

Here, we evaluate all possible combinations of a subset of chaperones in the human Hsp70 system and define the optimal chaperone set and optimal stoichiometry for luciferase disaggregation and reactivation. We establish that 115-7c stimulates the disaggregase activity of the optimized human Hsp70 system by ∼2-fold. We then report on a 115-7c analog that more potently stimulates the disaggregase activity of the human Hsp70 system against disordered luciferase aggregates and αSyn amyloid fibrils. The newly identified lead compound stimulates the disaggregase activity of multiple active human Hsp70, Hsp40, Hsp110 chaperone sets, with selectivity for combinations that include DnaJB1 or DnaJB4 as the Hsp40. We find that the relative stoichiometry of Hsp70, Hsp40, and Hsp110 dictates disaggregase activity. Remarkably, our lead compound shifts the composition of active chaperone stoichiometries by preferentially activating combinations with lower DnaJB1 concentrations. Collectively, our studies provide an important lead scaffold for further development via medicinal chemistry.

## Results

### Purification and activity of human Hsp70, Hsp40, and Hsp110 chaperones

We purified and tested the luciferase disaggregation and reactivation activity of two human Hsp70 proteins (Hsc70 [HspA8] and Hsp72 [HspA1A]), five human Hsp40 proteins (DnaJA1, DnaJA2, DnaJB1, DnaJB3, and DnaJB4), and two human Hsp110 proteins (Apg2 [HspH2] and Hsp105 [HspH1]) (Figure S1A-K). Hsc70 and Hsp72 are localized to the cytoplasm and nucleus and are found in the brain and many other tissues.^42,50^ Hsc70 is constitutively expressed whereas Hsp72 is a stress-inducible chaperone.^42,50,51^ DnaJA1, DnaJA2, DnaJB1 and DnaJB4 are localized to the cytoplasm and nucleus of many tissues, including the brain.^42^ DnaJB3 is primarily expressed in the testis and blood but is also expressed modestly in lung, spleen, blood, small intestine, heart, and kidney.^42,52^ Expression of DnaJB1 and DnaJB4 is heat-inducible, whereas the other Hsp40 proteins tested are not.^50^ Apg2 and Hsp105 are both constitutively localized to the cytoplasm and nucleus of multiple tissues, including brain.^42,50,53^ The Hsp105 isoform evaluated in this study is the α variant. There is also a smaller splice variant, Hsp105β, that is stress-inducible but is not tested here.^53^ Except for DnaJB3, all the chaperones tested here are expressed in brain, suggesting that they may cooperate to disaggregate proteins in neurons.^6,42,50^

The cDNA for each gene was cloned into the pE-SUMOpro expression vector and proteins were purified from *E. coli* via Ni-NTA and ion exchange chromatography. The purity of each protein ranges from ∼81%-98% (Figure S1A-J). We established that all the Hsp40 proteins significantly stimulate Hsc70 ATPase activity (Figure S2A). We also found that the Hsp70 proteins have intrinsic ATPase activity that is significantly stimulated by DnaJB1 (Figure S2B). Urea-denatured firefly luciferase forms a spectrum of aggregated species ∼500–2000 kDa and greater in size that are devoid of activity and very few luciferase species smaller than ∼400 kDa can be detected.^54^ We used this substrate to measure the protein disaggregation and reactivation ability of the Hsp70 system.^54^ We found that both Hsp110 proteins significantly stimulate luciferase disaggregation and reactivation by Hsc70 and DnaJB1 (Figure S2C). Thus, our purified Hsp70, Hsp40, and Hsp110 are all functional.

### Disaggregase activity of diverse three-component combinations of Hsp70, Hsp40, and Hsp110

We next assessed the disaggregase activity of all the possible three-component combinations of the purified Hsp70, Hsp40, and Hsp110 proteins. We designed an array-based, high-throughput version of the luciferase disaggregation and reactivation assay. We find that amongst the Hsp40 and Hsp110 proteins purified here, Hsc70 is generally more active than Hsp72 in disaggregating and reactivating luciferase (Figure 1A, B, S3A, B). Hsc70 is most active when paired with DnaJB4, less active when paired with DnaJB1, and even less active when paired with DnaJA2 (Figure 1A, S3A). Hsc70 displays very limited activity when paired with DnaJA1 or DnaJB3 (Figure 1A, S3A). Hsp72 is also most active when paired with DnaJB4 and less active when paired with DnaJB1 (Figure 1B, S3B). Hsp72 displays minimal activity when paired with DnaJA1, DnaJA2, or DnaJB3 (Figure 1B, S3B). DnaJB3 lacks the substrate-binding domains of DnaJB1 and DnaJB4 (Figure S1K), which may limit activity. Interestingly, three-component sets that contain DnaJB4 are significantly more active with Hsp105 as the Hsp110 component and less active with Apg2 (Figure 1A, B, S3A, B). Conversely, Hsc70 and DnaJA2 are equally active with Apg2 or Hsp105 (Figure 1A, S3A, B). Finally, three-component sets containing DnaJB1 have nearly equal activity regardless of the Hsp110 component, with Hsp105 being slightly less active (Figure 1A, B, S3A, B). Thus, we define a range of activities and productive interactions among human Hsp70, Hsp40, and Hsp110 proteins for disaggregation and reactivation of chemically denatured luciferase.

**Figure 1.**
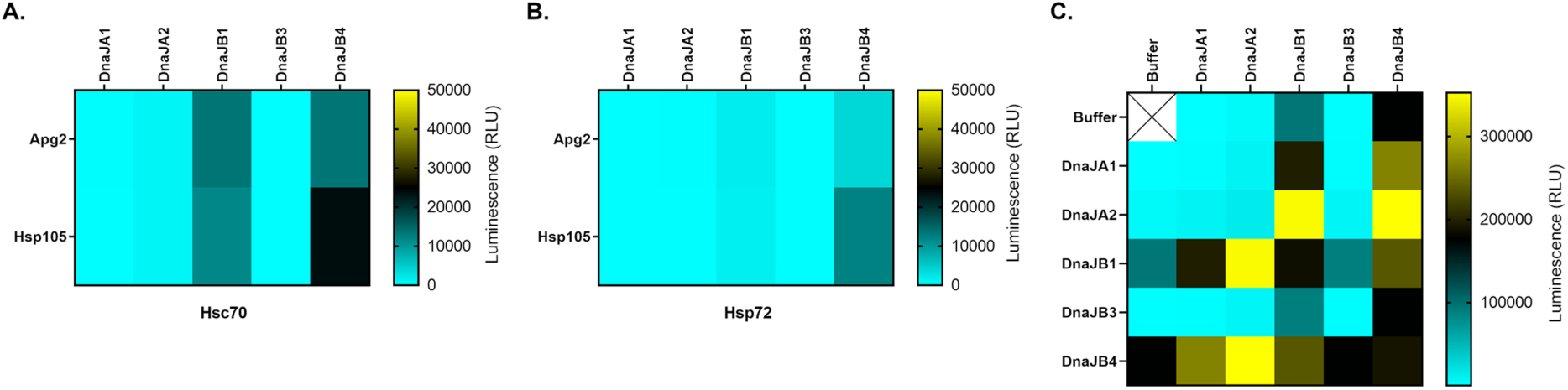
Distinct combinations of human Hsp70, Hsp40, and Hsp110 display diverse levels of protein-disaggregase activity. **(A)** Heat map showing the luciferase disaggregase and reactivation activity of Hsc70 with every pairwise combination of the Hsp40 (DnaJA1, DnaJA2, DnaJB1, DnaJB3, or DnaJB4) and Hsp110 (Apg2 or Hsp105) proteins purified. 0.4μM Hsc70, 0.2μM Hsp40, and 0.04μM Hsp110 were combined with 100nM luciferase aggregates (monomeric concentration), 1% DMSO, and an ATP-regenerating system. Colors represent mean luminescence (n=4). **(B)** Heat map showing the luciferase disaggregase and reactivation activity of Hsp72 with every pairwise combination of the Hsp40 and Hsp110 proteins purified. 0.4μM Hsp72, 0.2μM Hsp40, and 0.04μM Hsp110 were combined with 100nM luciferase aggregates (monomeric concentration), 1% DMSO, and an ATP-regenerating system. Colors represent mean luminescence (n=3). **(C)** Heat map showing the luciferase disaggregase and reactivation activity of Hsc70 and Apg2 with pairwise combinations of the Hsp40 proteins. 1μM Hsc70, 0.1μM Apg2, and 0.25μM of each Hsp40 were combined with 100nM luciferase aggregates (monomeric concentration) and an ATP-regenerating system. The column and row labeled buffer have 0.25μM of a single Hsp40. The buffer vs. buffer condition (i.e., no Hsp40) was not determined and is indicated by a crossed out white box. Boxes along the diagonal have 0.5μM of a single Hsp40. Data are symmetric across the diagonal. Colors represent mean luminescence (n=3-6). See also **Figure S1, S2, and S3**.

### Class A and class B Hsp40 proteins can synergize to yield enhanced disaggregase activity

Prior studies have suggested that class A and class B Hsp40 proteins can synergize to promote greater protein disaggregase activity with Hsp70 and Hsp110 than if only one class A or class B Hsp40 is used.^15^ Specifically, Hsc70 and Apg2 combined with either DnaJA2 or DnaJB1 were found to have modest luciferase disaggregase and reactivation activity, but when combined together Hsc70, Apg2, DnaJA2, and DnaJB1 showed increased disaggregase and reactivation activity at the same total Hsp40 concentration.^15^

We next determined whether other such synergistic pairs of Hsp40 proteins might exist. We used the array-based luciferase disaggregation and reactivation assay to test pairwise combination of DnaJA1, DnaJA2, DnaJB1, DnaJB3, and DnaJB4 with Hsc70 and Apg2. DnaJB1 (0.25μM) enables modest disaggregase and reactivation activity as the sole Hsp40 with Hsc70 (1.0μM) and Apg2 (0.1µM), but when either DnaJA1 (0.25μM) or DnaJA2 (0.25μM) is added we find a marked increase in disaggregase and reactivation activity (Figure 1C, S3C, D, E). DnaJA1 or DnaJA2 have limited activity as the sole Hsp40 component or when combined, indicating that these class A Hsp40s synergize with DnaJB1 (Figure 1C, S3C, D, E). Indeed, combining DnaJA1 with DnaJB1 increased activity by ∼2-fold over the predicted additive effect (Figure 1C, S3C, E), whereas combining DnaJA2 with DnaJB1 increased activity by ∼3.6-fold over the predicted additive effect (Figure 1C, S3D, E). Thus, DnaJA1 or DnaJA2 synergize with DnaJB1 to promote luciferase disaggregation and reactivation.

By contrast, DnaJB3 facilitated minimal disaggregase and reactivation activity as the sole Hsp40 with Hsc70 and Apg2 (Figure 1C, S3F). When DnaJB3 is combined with DnaJA1 or DnaJA2 there is also very little activity (Figure 1C, S3C, D, F). Thus, DnaJA1 or DnaJA2 do not synergize with DnaJB3 to promote luciferase disaggregation and reactivation.

DnaJB4 elicits strong disaggregase and reactivation activity as the sole Hsp40 with Hsc70 and Apg2, but when either DnaJA1 or DnaJA2 is added we observe markedly increased disaggregase and reactivation activity (Figure 1C, S3G). Indeed, combining DnaJA1 with DnaJB4 increased activity by ∼1.6-fold over the predicted additive effect (Figure 1C, S3C, G), whereas combining DnaJA2 with DnaJB4 increased activity by ∼2-fold over the predicted additive effect (Figure 1C, S3D, G). Thus, DnaJA1 or DnaJA2 synergize with DnaJB4 to promote luciferase disaggregation and reactivation.

Interestingly, combining DnaJB1 (0.25μM) and DnaJB4 (0.25μM) also shows greater disaggregase and reactivation activity, but the increase is nearly equal to the sum of the activity of the two Hsp40 proteins separately (Figure 1C, S3E, G). By contrast, combining DnaJB3 with DnaJB1 or DnaJB4 had no effect on activity (Figure 1C, S3E, F, G). Overall, these data reveal that class A and class B Hsp40 proteins can, but do not always (e.g., DnaJA1 and DnaJB3), synergize to yield greater disaggregase and reactivation activity.^15,29,43^ They also suggest that pairs of class A or pairs of class B Hsp40s lack synergistic effects in luciferase disaggregation and reactivation.

### Analogs of a small molecule Hsp70 agonist further stimulate disaggregation and reactivation of luciferase

The dihydropyrimidine, 115-7c (Figure S4), enhances the luciferase refolding activity of the homologous bacterial system composed of DnaK, DnaJ, and GrpE and a derivative enhances single-turnover ATP hydrolysis of human Hsp70.^46,48,49^ However, it is unknown if 115-7c stimulates the human Hsp70 chaperone system to disaggregate and reactivate luciferase trapped in larger aggregated species. 115-7c can reduce Htt-polyQ aggregation in HEK293T cells and reduce αSyn aggregation in H4 neuroglioma cells, but it is unclear if these effects are directly due to disaggregation of the disease protein.^46,55,56^ Therefore, we next determined whether 115-7c or structurally related analogs could enhance the disaggregase activity of the human Hsp70 system in biochemical assays. We used Hsc70, DnaJB1, and Apg2 to test for small-molecule stimulation because this chaperone set is found in the brain and disaggregates human disease-related protein aggregates including αSyn, Htt-polyQ, and tau.^19–22^ Hsc70, DnaJB1, and Apg2 also show robust luciferase disaggregase and reactivation activity, allowing for the identification of small-molecule enhancers using a highly scalable assay. We determined that chaperone concentrations that yield ∼10% of maximal effect (EC_10_) are Hsc70 (0.4µM), DnaJB1 (0.2µM), and Apg2 (0.04µM). Testing drug-like small molecules at the EC_10_ allows for a large dynamic range to identify stimulators of Hsp70, Hsp40, and Hsp110 disaggregase activity.

We found that 115-7c (25μM) enhances the disaggregase activity of the Hsp70 system by ∼2-fold over the solvent (DMSO) control (Figure 2A). Next, we tested structural analogs of 115-7c to uncover more potent stimulators of Hsp70 disaggregase activity (Figure S4). These compounds were synthesized according to published procedures.^46,57^ Compounds 8, 16, 17, 18, and 19 enhanced disaggregase and reactivation activity significantly over the solvent (DMSO) control (Figure 2A). Among these compounds, 18 was the most potent stimulator with a ∼7-fold increase in disaggregase and reactivation activity over DMSO and was the only analog to stimulate activity significantly better than 115-7c (Figure 2B). Compounds 16, 17, 18, and 19 all share the same core structure as 115-7c and differ in the functional groups in the substituents. Furthermore, these compounds share an ester functional group added to the 115-7c carboxylate (Figure 2C, red). Compound 8 has a naphthalene substituent rather than a dichlorophenyl group (Figure 2C, blue). Compound 8 also has two extra methylene groups in the carbon chain leading to the carboxylic acid (Figure 2C, red). Compound 7 bears a naphthalene substituent but does not significantly stimulate disaggregase activity compared to DMSO (Figure 2A, S4).

**Figure 2.**
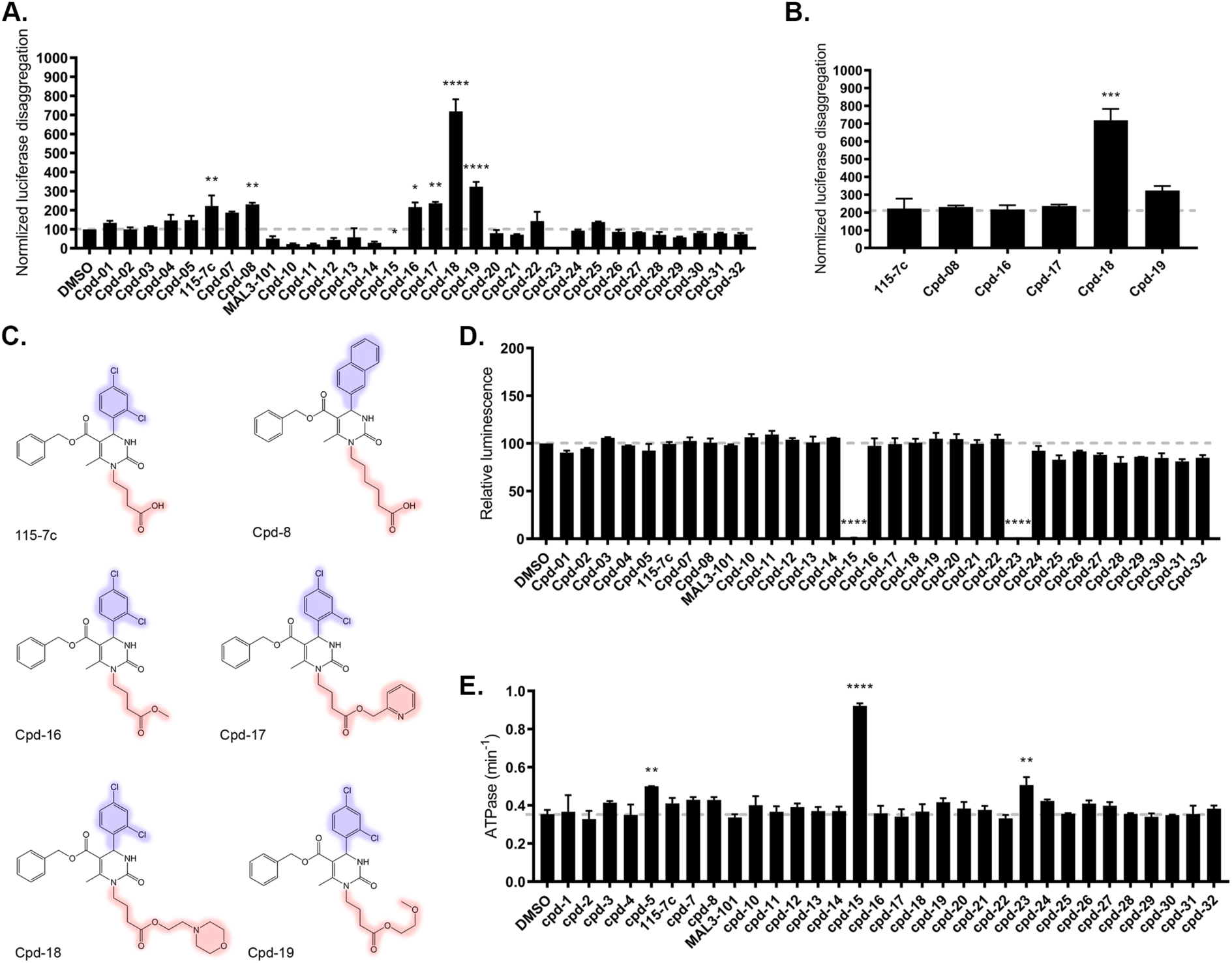
Dihydropyrimidine 115-7c and structural analogs stimulate the luciferase disaggregation and reactivation activity of Hsc70, DnaJB1, and Apg2. **(A)** Luciferase disaggregation and reactivation activity of Hsc70, DnaJB1, and Apg2 in the presence of 1% DMSO or 25μM compound (final 1% DMSO). 0.4μM Hsc70, 0.2μM DnaJB1, and 0.04μM Apg2 were combined with 100nM luciferase aggregates (monomeric concentration), an ATP-regenerating system, and either DMSO or compound. Values are normalized to DMSO treated control and are means ± SEM (n=2). Data were analyzed using one-way ANOVA followed by Dunnett’s MCT compared to DMSO control (*p < 0.05, **p < 0.01, ****p < 0.0001). **(B)** Data from (A) were taken for the active compounds 115-7c, 8, 16, 17, 18, and 19 for statistical analysis. Values are normalized to DMSO treated control and are means ± SEM (n=2). Data were analyzed using one-way ANOVA followed by Dunnett’s MCT compared to 115-7c (***p < 0.001). **(C)** Chemical structures of 115-7c and active analogs. Structural differences between compounds highlighted in red and blue. **(D)** Activity of native luciferase in the presence of 1% DMSO or 25μM compound (final 1% DMSO). 16nM native luciferase was combined with an ATP-regenerating system and either DMSO or compound. Values are normalized to DMSO treated control and are means ± SEM (n=2). Data were analyzed using one-way ANOVA followed by Dunnett’s MCT compared to DMSO control (****p < 0.0001). **(E)** ATPase activity of Hsc70, DnaJB1, and Apg2 in the presence of 1% DMSO or 25μM compound (final 1% DMSO). 0.4μM Hsc70, 0.2μM DnaJB1, and 0.04μM Apg2 were incubated with 1mM ATP. Values represent means ± SEM (n=2). Data were analyzed using one-way ANOVA followed by Dunnett’s MCT compared to DMSO control (**p < 0.01, ****p < 0.0001). See also **Figure S4** and **S5**.

We next tested the effects of these compounds on native luciferase in the absence of any chaperones. This assay would reveal any compounds that directly affect the activity of the reactivated luciferase instead of chaperone-mediated disaggregation and reactivation activity. Compounds 8, 16, 17, 18, and 19 do not enhance the activity of native luciferase (Figure 2D), strongly suggesting that the increase in the luminescence signal arises from enhanced disaggregase and reactivation activity (Figure 2A). We also observed that compounds 15 and 23 inhibit native luciferase, which likely explains why little luciferase activity was recovered by the Hsp70-disaggregase system in the presence of these compounds (Figure 2A, S4).^58,59^

### Analogs of a small-molecule Hsp70 agonist do not stimulate the ATPase activity of the human Hsp70-disaggregase system

We next assessed whether the compounds modulate the ATPase activity of the Hsp70-disaggregase system. The nucleotide state of Hsp70 determines both its structural conformation and its affinity for substrates, as Hsp70 uses ATP hydrolysis to regulate the capture and release of its substrates.^29^ Compounds 115-7c, 8, 16, 17, 18, and 19 did not stimulate global steady state ATPase activity of Hsc70, DnaJB1, and Apg2 (Figure 2E). This finding suggests that the stimulation of disaggregase activity (Figure 2A) is not due to enhanced global ATPase activity. Thus, we propose that stimulation of disaggregase activity arises from improved efficiency of ATP utilization, i.e., ATP hydrolysis is more likely to be coupled to productive disaggregation.

At first glance, the lack of ATPase stimulation might appear unexpected since 115-7c binds at the Hsp70 and Hsp40 interface, inducing an allosteric change in the ATP-binding site, which is mediated by an amino-acid-relay system, and ultimately promotes Hsp40-stimulated ATPase activity^49^. However, stimulation of ATPase activity by 115-7c in single-turnover assays was previously reported for bacterial chaperones DnaK, DnaJ, and GrpE, the yeast chaperones Ssa1 and Ydj1, and human Hsc70 with bacterial DnaJ.^46,49,60^ None of these experiments included optimized assays with Hsp110. Here, we report the effects of these compounds on the ATPase activity of the human Hsp70-Hsp40-Hsp110 system, which has not been examined before.

In contrast to the effects of compounds 115-7c, 8, 16, 17, 18, and 19, compounds 5, 15, and 23 stimulate ATPase activity (Figure 2E). Compound 5 has a bulkier and more rigid carboxylic tail than 115-7c (Figure S4). Compounds 15 and 23 both contain a tetrazole bioisostere in lieu of the carboxylate (Figure S4), but they abolish native luciferase activity and so their effects on luciferase disaggregase and reactivation activity could not be determined (Figure 2D). Thus, the tetrazole ring may enable the stimulation of ATPase activity, the inhibition of luciferase activity, or both. This result is somewhat unexpected as compounds 15 and 23 were originally designed to selectively inhibit the Simian Virus protein T-antigen.^59^ However, both compounds were found to lack selectivity and also inhibit the ATPase activity of human Hsp70 and Hsp40 in the absence of Hsp110.^59^ Notably, in our studies, Hsp110 is included, which may directly contribute to ATPase activity (i.e., Hsp110 has intrinsic ATPase activity^13^), regulate the ATPase activity of Hsp70, or both.

### Structure-activity relationship of 115-7c derivatives and stimulation of the human Hsp70-disaggregase system

The parent scaffold 115-7c exhibits a ∼2-fold increase in the ability of the Hsp70 system to disaggregate and reactivate luciferase (Figure 2A). Compounds 1, 2, 3, and 4 vary the dichlorobenzyl moiety by either removing one of the chloride atoms or replacing a chloride with a fluoride or a trifluoromethyl group (Figure S4). These modifications result in a loss of the statistically significant stimulation observed with 115-7c (Figure 2A). The benzoic acid modification in compound 5 increases ATPase activity but does not significantly increase disaggregase and reactivation activity (Figure 2A, E). Analog 116-9e (not tested here), where the dichlorobenzyl group is replaced with a more extended biphenyl moiety,^57^ inhibits the interaction between DnaK and DnaJ, suggesting that this region of the molecule is important for activity.^49^ In contrast, in compounds 7 and 8 the dichlorobenzyl moiety is replaced with a bulkier naphthalene ring (Figure S4). Compound 7 is less active than 115-7c, which suggests that the bulkier naphthalene moiety in compound 7 reduces its interaction with Hsp70 and Hsp40 in a similar manner to the biphenyl moiety in compound 116-9e. Compound 8 retains similar activity to 115-7c despite having the same naphthalene moiety as compound 7. Therefore, the longer and more flexible carbon chain ending in a carboxylic acid in compound 8 may compensate for the negative effects of the naphthalene substituent.

MAL3-101 is a small-molecule inhibitor of J-domain stimulated yeast Hsp70 ATPase activity.^45,61^ Compounds 10, 11, 12, 13, and 14 share structural homology to MAL3-101 (Figure S4).^57^ MAL3-101 and 115-7c also share the same pyrimidine core, but MAL3-101 has an expanded number of functional groups, is chemically more complex, and is approximately twice the molecular weight of 115-7c (Figure S4). We find that MAL3-101 along with compounds 10 through 14 do not inhibit global ATPase activity of the human Hsp70 system but they do inhibit the disaggregase and reactivation activity (Figure 2A, E). Compounds 15 and 23 both have the dicholorobenzyl moiety replaced with a bulkier biphenyl moiety, and they have a tetrazole ring instead of a carboxylate (Figure S4). Compounds 15 and 23 stimulate the ATPase activity of the Hsp70 system, but inhibit native luciferase (Figure 2D, E). Thus, their effect on luciferase disaggregation and reactivation by the Hsp70 system could not be determined.

Compounds 16 through 22 all share the same scaffold as 115-7c but differ by additional ester groups added to the carboxylate tail (Figure S4).^46^ Compound 18 is the only scaffold that stimulates disaggregase and reactivation activity significantly more than 115-7c (Figure 2B). Compound 18 has a flexible side chain terminating in a morpholino group (Figure 2C, S4). Compound 19 instead has a methoxyethyl group attached to the carboxylate and is not as active as compound 18 (Figure 2A, C, S4). By contrast, the methyl ester derivative, compound 16, exhibits similar activity to 115-7c (Figure 2A, C, S4), suggesting that a small group is well tolerated at this site. Compound 17 has a 2-pyridyl ester and lacks the enhanced activity of compound 18, suggesting that the oxygen atom or the ring flexibility in the morpholino group is important for activity (Figure 2A, C, S4). Like compound 17, compounds 20 and 21 have larger hydrophobic ester substituents that prevent stimulation of disaggregase activity (Figure 2A, S4). In turn, compound 22 has a bulky quinolone that is also related to compound 17 and prevents significant stimulation of disaggregase activity (Figure 2A, S4). Compound 26 has a cyano methyl ester and does not show activity in stimulating luciferase disaggregation or ATPase activity, which is interesting given that this functional group is only slightly larger than the methyl ester derivative in compound 16 (Figure 2A, S4).^46^ Compound 26 reduces αSyn aggregation in H4 neuroglioma cells.^46^ However, given our results, this cellular activity might not be due to direct stimulation of the Hsp70-disaggregase system, but rather compound 26 is converted to an acid, i.e., compound 115-7c, in the cell.^46^

Another cohort of molecules include compounds 24 and 29, which contain fused tetrahydropyridimines and do not significantly affect disaggregase or ATPase activity (Figure 2A, E, S4).^62^ Analogs 27, 28, 30, and 32 contain the dihydropyrimidine core but are smaller and structurally less complex than 115-7c and also do not significantly affect disaggregase or ATPase activity (Figure 2A, E, S4). Notably, compounds 25 and 31 lie structurally in between 115-7c and MAL3-101 (Figure S4) as they have multiple functional groups attached to the carboxylate tail, thereby forming an amide bond rather than the ester in compound 18 (Figure S4). Accordingly, they more closely resemble MAL3-101, and neither compound 25 nor 31 affect ATPase or disaggregase activity (Figure 2A, E, S4).

Overall, compounds 8, 16, 17, 18, and 19 exhibit similar activity to 115-7c and significantly enhance the activity of the human Hsp70-disaggregase system (Figure 2A). However, only compound 18 shows a significantly enhanced stimulation of disaggregase activity when compared to 115-7c (Figure 2B). Our results suggest that certain functional groups added to the carboxylic tail might make key contacts with Hsp70 and thereby contribute to the effect of compound 18 (Figure 2C, red).

We next assessed whether the small-molecule stimulators, 115-7c, 8, 16, 17, 18, and 19, exhibit drug-like character and calculated their physicochemical properties using SwissADME (Table S1).^63^ With the exception of compounds 17, 18, and 19, which have M_W_’s of 535-590, the compounds pass Lipinski’s rule of five (Table S1),^64^ which distinguishes a large majority of known FDA-approved oral drugs and predicts satisfactory permeability and absorption.^65^ In general, an orally active drug has no more than one violation of the following rules: (1) the molecule has no more than five H-bond donors (HBD); (2) the molecule has no more than 10 H-bond acceptors (HBA); (3) the molecular weight (M_W_) is <500 Da; and (4) cLogP is < 5.^64^ However, the M_W_ rule is the most commonly violated rule in FDA-approved drugs,^65^ indicating that high M_W_ may be a more tolerable physical property.^66,67^ However, none of the compounds were predicted to be blood-brain-barrier permeable, which will be vital to address via medicinal chemistry in the future.

### Compound 18 stimulates luciferase disaggregation and reactivation by the human Hsp70 system in a dose-dependent manner

We focused on compound 18 since it shows the greatest stimulation of the human Hsp70-disaggregase system among the 115-7c analogs. We treated Hsc70 (0.4µM), DnaJB1 (0.2µM), and Apg2 (0.04µM) with compound 18 (or the DMSO control) at a range of concentrations. Compound 18 stimulated luciferase disaggregase and reactivation by the Hsp70-disaggregase system in a dose-dependent manner but reaches a maximum at 25μM and stimulation declined at 100µM (Figure 3A, blue). We found the EC_50_ (half maximal effective concentration) of compound 18 was ∼9.2±1.9μM. Notably, none of the tested concentrations of compound 18 affected native luciferase activity (Figure 3A, red). Thus, the decline in Hsp70-reactivated luciferase activity at high concentrations of compound 18 is not a result of direct inhibition of luciferase by compound 18. Moreover, compound 18 does not cause direct disaggregation or reactivation of luciferase aggregates in the absence of the chaperones (Figure S5A).

**Figure 3.**
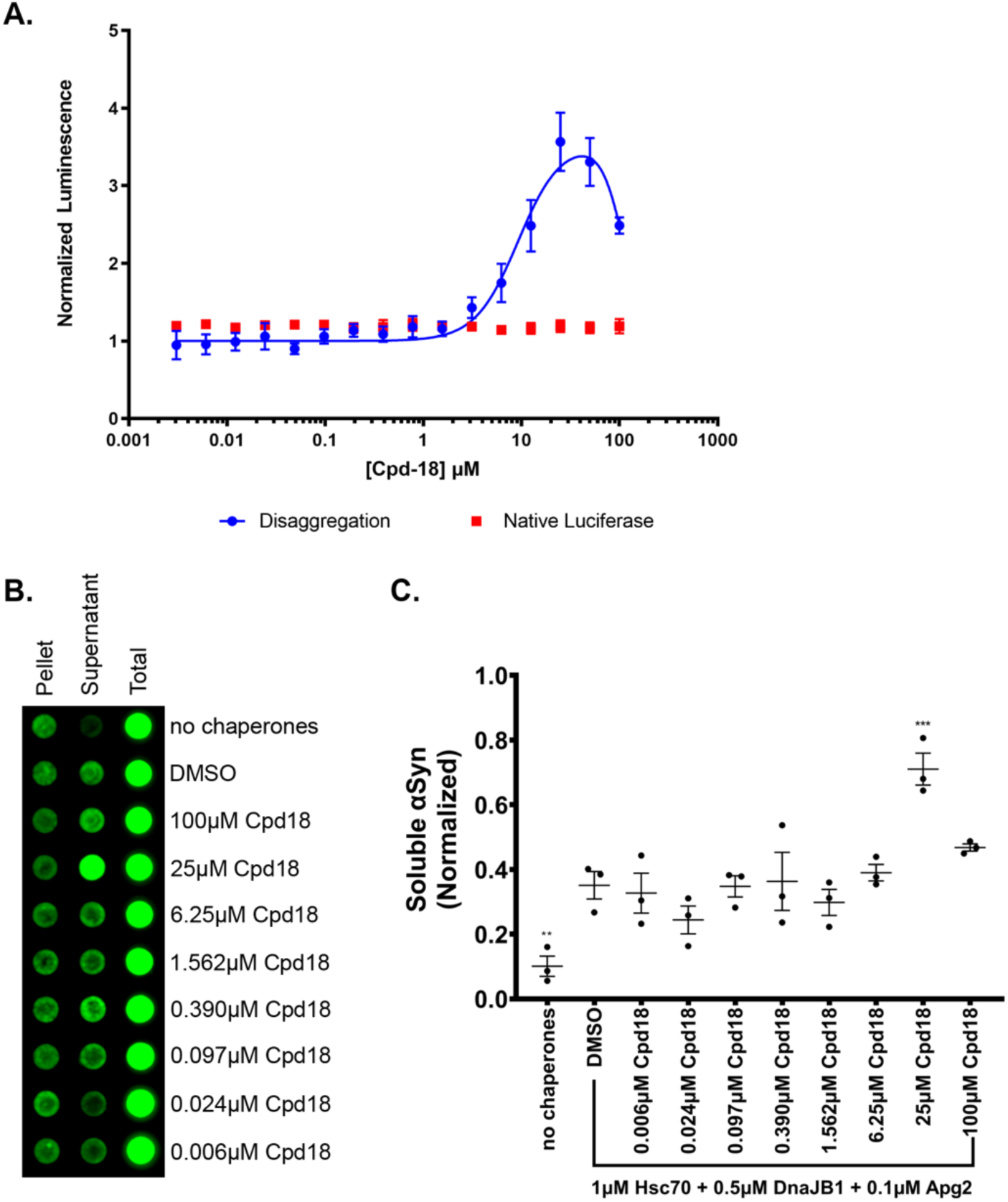
Compound 18 stimulates the luciferase and αSyn disaggregase activity of Hsc70, DnaJB1, and Apg2 in a dose-dependent manner. **(A)** In blue, luciferase disaggregation and reactivation activity of Hsc70, DnaJB1, and Apg2 in the presence of 1% DMSO or 0.003-100μM compound 18 (final 1% DMSO). 0.4μM Hsc70, 0.2μM DnaJB1, and 0.04μM Apg2 were combined with 100nM luciferase aggregates (monomeric concentration), an ATP-regenerating system, and either DMSO or compound. Values are normalized to DMSO treated control and are means ± SEM (n=3). Nonlinear curve fitting was performed with GraphPad Prism using the bell-shaped dose response curve fitting. In red, activity of native luciferase in the presence of 1% DMSO or 0.003-100μM compound 18 (final 1% DMSO). 16nM native luciferase was combined with an ATP-regenerating system and either DMSO or compound. Values are normalized to DMSO treated control and are means ± SEM (n=3). **(B)** Representative dot blot showing αSyn content in pellet, supernatant, and total fractions after 0.5μM αSyn PFFs were treated with 1μM Hsc70, 0.5μM DnaJB1, and 0.1μM Apg2 at 37°C while shaking at 300rpm for 90min. Each row was treated with the indicated concentration of compound 18 with a final concentration of 1% DMSO. 10% of the total reaction, supernatant, or resuspended pellet were loaded onto the blot and stained with SYN211. **(C)** Quantification of three trials of the αSyn disaggregation assay described in (B). Dot blots were quantified using FIJI integrated density measurements. Soluble αSyn in the supernatant fraction was normalized by dividing by the total loaded αSyn for each corresponding condition and then plotted in GraphPad Prism. Y-axis represents the normalized soluble αSyn calculated. Individual data points shown as dots, bars represent mean ± SEM (n=3). Data were analyzed using one-way ANOVA followed by Dunnett’s MCT compared to DMSO with chaperones control (**p < 0.01, ***p < 0.001). See also **Figure S5**.

### Compound 18 can stimulate disaggregation of α-Syn amyloid fibrils by Hsc70, DnaJB1, and Apg2

Luciferase forms amorphous aggregates that lack the ordered amyloid structure observed in proteins that aggregate in neurodegenerative disease, including αSyn, amyloid-beta, and tau.^6,7^ To determine if compound 18 stimulates the disaggregase activity of the Hsp70 system against ordered amyloid aggregates, we measured αSyn preformed fibril (PFF) disaggregation. These αSyn PFFs are reactive to the amyloid dye thioflavin-T and induce a PD-like phenotype in mice.^68^ Previous studies have shown that αSyn can be disaggregated by Hsc70, DnaJB1, and Apg2 and that αSyn fibrils are preferentially dissembled from the ends into monomers, but are not fragmented.^19,20,24,26,28^ Thus, we treated αSyn PFFs (0.5μM monomer) with Hsc70 (1µM), DnaJB1 (0.5µM), and Apg2 (0.1µM) in the presence or absence of compound 18 at a range of concentrations. Then, the products were centrifuged and separated into supernatant and pellet fractions. The αSyn content of the supernatant, pellet, and total fractions was measured by dot blot using anti-SYN211 antibody (Figure 3B).

When we treated αSyn PFFs with chaperones and DMSO, αSyn was disaggregated from the insoluble fibrils and released into the supernatant (Figure 3B). Disaggregase activity was further stimulated by compound 18 with up to a ∼2-fold increase over the DMSO control (Figure 3C). Notably, enhanced disaggregase activity was only statistically significant at a final concentration of 25μM compound 18 (Figure 3C). Furthermore, compound 18 does not cause direct disaggregation of αSyn PFFs (Figure S5B, C). We conclude that compound 18 stimulates the disaggregase activity of the Hsp70 system against both aggregated luciferase and αSyn PFFs *in vitro*, suggesting that the mechanism of disaggregation for amorphous and ordered aggregates share common mechanistic steps.

### Compound 18 does not stimulate luciferase disaggregation and reactivation by AAA+ disaggregases

To ensure compound 18 was specific for Hsp70, Hsp40, and Hsp110, we next assessed whether it stimulates luciferase disaggregation and reactivation by AAA+ disaggregases, which bear no resemblance to the Hsp70-disaggregase system.^69^ We selected the potentiated Hsp104 variant, Hsp104^K358D,70^ or the human mitochondrial protein disaggregase Skd3 (we used the PARL-activated form of Skd3, _PARL_Skd3).^71–73^ Neither Hsp104^K358D^ nor _PARL_Skd3 require Hsp70, Hsp40, or Hsp110 to disaggregate and reactivate luciferase.^70,72^ Thus, we can ask whether compound 18 stimulates luciferase disaggregation and reactivation by diverse protein disaggregases or whether this activity is specific for Hsp70, Hsp40, and Hsp110. Compound 18 did not stimulate luciferase disaggregation and reactivation by Hsp104^K358D^ or _PARL_Skd3 (Figure S5D). Thus, compound 18 displays a selective ability to stimulate luciferase disaggregation and reactivation by Hsp70, Hsp40, and Hsp110.

### Compound 18 stimulates Hsp70, Hsp40, Hsp110 chaperone sets containing DnaJB1 or DnaJB4

Compound 18 was unable to stimulate disaggregase activity by AAA+ disaggregases, but we wondered whether it could stimulate disaggregase activity of diverse sets of human Hsp70, Hsp40, and Hsp110 chaperones. Therefore, we tested all possible three component combinations of the two Hsp70s, five Hsp40s, and two Hsp110s in the luciferase disaggregation and reactivation assay. Ultimately, we found that compound 18 significantly increased the disaggregase activity of Hsc70 and DnaJB1 or DnaJB4 with either of the Hsp110 proteins (Figure 4A, B). By contrast, compound 18 failed to stimulate the disaggregase activity of Hsc70 and DnaJA2 with either of the Hsp110 proteins (Figure 4A, B). Furthermore, Hsc70 with DnaJA1 or DnaJB3 is inactive with either of the Hsp110 proteins regardless of whether compound 18 was present (Figure 4A, B). A similar trend was evident with Hsp72. Compound 18 stimulated the activity of Hsp72 and DnaJB1 or DnaJB4 with either of the Hsp110 proteins (Figure 4C, D) and failed to stimulate Hsp72 and DnaJ1, DnaJA2, or DnaJB3 with either Apg2 or Hsp105 (Figure 4C, D). Thus, compound 18 is unable to stimulate the disaggregase activity of any combination of Hsp70, Hsp40, and Hsp110.

**Figure 4.**
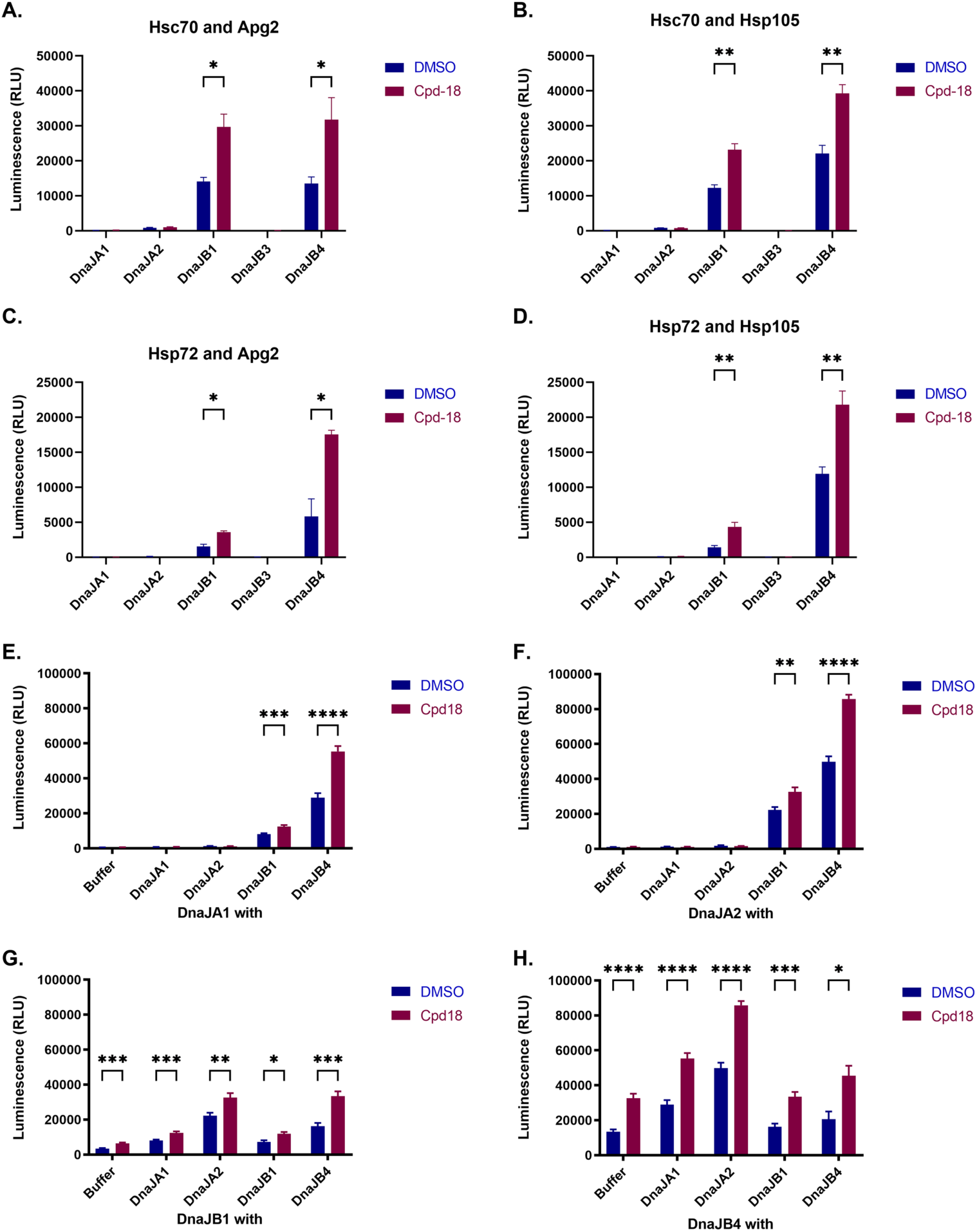
Compound 18 is not specific for Hsc70, DnaJB1, and Apg2 and stimulates the activity of multiple Hsp70, Hsp40, Hsp110 chaperone sets. **(A-D)** Bar graphs showing the luciferase disaggregase and reactivation activity of (A) Hsc70 and Apg2, (B) Hsc70 and Hsp105, (C) Hsp72 and Apg2, or (D) Hsp72 and Hsp105 with DnaJA1, DnaJA2, DnaJB1, DnaJB3, or DnaJB4. Each chaperone set was tested with 1% DMSO (blue) or 25μM compound 18 (red). 0.4μM Hsc70, 0.2μM Hsp40, and 0.04μM Hsp110 were combined with 100nM luciferase aggregates (monomeric concentration), final 1% DMSO, and an ATP-regenerating system. Values represent mean ± SEM (n=3-6). Data were analyzed using unpaired t-test for each chaperone set comparing DMSO treated with compound 18 treated (*p < 0.05, **p<0.01). **(E-H)** Bar graphs showing the luciferase disaggregase and reactivation activity of pairwise combinations of (E) 0.1µM DnaJA1, (F) 0.1µM DnaJA2, (G) 0.1µM DnaJB1, (H) 0.1µM DnaJB4 plus either buffer, 0.1µM DnaJA1, 0.1µM DnaJA2, 0.1µM DnaJB1, or 0.1µM DnaJB4 with Hsc70 and Apg2. Each chaperone set was tested with 1% DMSO (blue) or 25μM compound 18 (red). 0.4μM Hsc70, 0.2μM Hsp40 (except for the buffer control, which has 0.1µM Hsp40), and 0.04μM Hsp110 were combined with 100nM luciferase aggregates (monomeric concentration), final 1% DMSO, and an ATP-regenerating system. Values represent mean ± SEM (n=3-10). Data were analyzed using unpaired t-test for each chaperone set comparing DMSO treated with compound 18 treated (*p < 0.05).

When we added compound 18 to four-component chaperone sets comprising Hsc70, Apg2, and two Hsp40 proteins, a similar result was observed. Specifically, reactions containing either DnaJB1 or DnaJB4 are stimulated whereas sets that lack a class B Hsp40 (i.e., those composed of only DnaJA1, DnaJA2, or both) are not stimulated (Figure 4E-H). Compound 18 does not stimulate Hsc70, DnaJA2, and Apg2 (Figure 4F). Interestingly, however, when DnaJA2 is paired with DnaJB1 or DnaJB4 the disaggregase activity is greatly enhanced above DnaJB1 or DnaJB4 alone (Figure 4F, G, H), and this activity is further stimulated by compound 18 (Figure 4F). Thus, only four-component chaperone sets that contain DnaJB1 or DnaJB4 are stimulated by compound 18.

In general, compound 18 is not specific for Hsc70, DnaJB1, and Apg2, and stimulates most of the active chaperone sets including combinations of Hsc70 and Hsp72, DnaJB1 and DnaJB4, and either of the Hsp110 proteins (Figure 4). Hsc70 and DnaJA2 are marginally active with either of the Hsp110 proteins, but these chaperone sets are not stimulated by compound 18. This finding suggests that the effect of compound 18 is specific to DnaJB1/4-containing chaperone sets. In addition, the selectivity of compound 18 is primarily defined by the Hsp40 component since for any given Hsp40, stimulation is all-or-none for Hsp70 and Hsp110 combinations (Figure 4A-D). The compound selectively promotes stimulation of Hsp70, Hsp40, and Hsp110 sets containing DnaJB1 and DnaJB4, including in the presence of DnaJA1 or DnaJA2, and stimulation is largely independent of the identity of the Hsp70 and Hsp110 components tested here.

### Compound 18 stimulates the Hsp70-disaggregase system at diverse chaperone stoichiometries

The activity of the Hsp70-disaggregase system is highly sensitive to the stoichiometric composition of the chaperone components, and prior studies have reported different stoichiometric ratios can be operational for disaggregase activity^13,16,18,20–22,27^. Therefore, we next systematically explored a landscape of Hsc70, DnaJB1, and Apg2 stoichiometries for luciferase disaggregase and reactivation activity (Figure 5). We kept the Hsc70 concentration constant (1µM) and varied the concentration of DnaJB1 (0-10µM), as well as the concentration of Apg2 (0-1µM) and thereby explored a matrix of 48 different chaperone stoichiometries. Under these conditions, the optimal chaperone concentrations are 1μM Hsc70, 5μM DnaJB1, and 0.1μM Apg2 (Figure 5A, C). Interestingly, disaggregase and reactivation activity does not correlate monotonically with DnaJB1 or Apg2 concentrations. Apg2 is required for robust disaggregase activity,^16^ but higher Apg2 concentrations can inhibit disaggregase activity (Figure 5A, C). In particular, optimal disaggregase and reactivation activity is observed at more moderate levels of Apg2 (i.e., 0.05μM and 0.1μM; Figure 5A, C). DnaJB1 is also required for robust disaggregase activity,^16^ but at low concentrations of Apg2 such as 0.005μM, excess DnaJB1 inhibits disaggregase and reactivation activity (Figure 5A, C).

**Figure 5.**
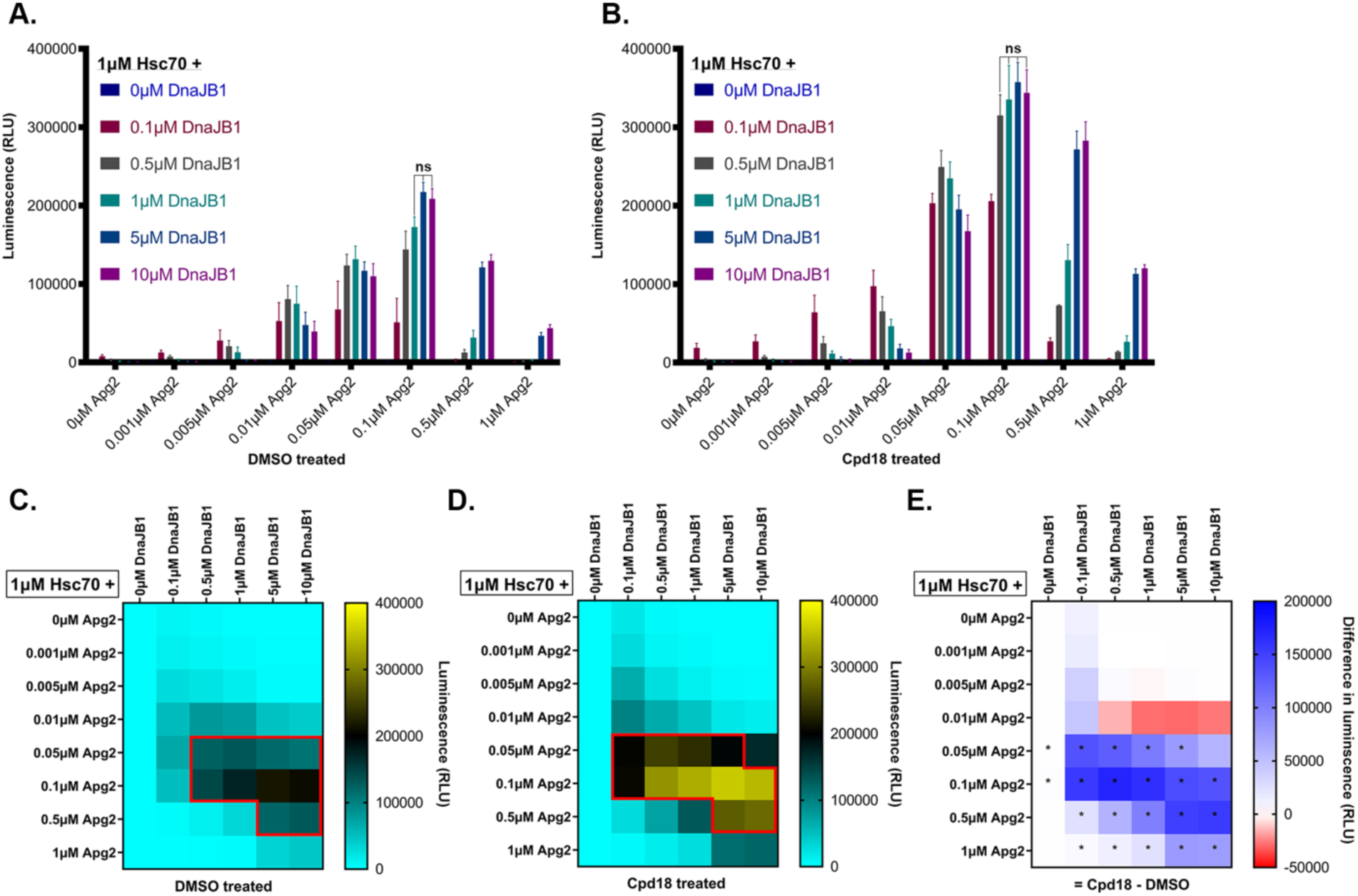
Compound 18 stimulates the Hsp70-disaggregase system in diverse positions within the landscape of chaperone stoichiometries. **(A)** Luciferase disaggregase and reactivation activity of Hsc70, DnaJB1, and Apg2 at a range of stoichiometries treated with DMSO. 1μM Hsc70, 0μM-10μM DnaJB1, and 0μM-1μM Apg2 were combined with 100nM luciferase aggregates (monomeric concentration), an ATP-regenerating system, and final 1% DMSO. Values are means ± SEM (n=3). Data were analyzed using one-way ANOVA followed by Dunnett’s MCT compared to the optimal stoichiometry: 1μM Hsc70, 5μM DnaJB1, and 0.1μM Apg2 (ns = p>0.05, all other values have p<0.05). **(B)** Luciferase disaggregase and reactivation activity of Hsc70, DnaJB1, and Apg2 at a range of stoichiometries treated with compound 18. 1μM Hsc70, 0-10μM DnaJB1, and 0-1μM Apg2 were combined with 100nM luciferase aggregates (monomeric concentration), an ATP-regenerating system, and 25μM compound 18 (final 1% DMSO). Values are means ± SEM (n=3). Data were analyzed using one-way ANOVA followed by Dunnett’s MCT compared to the optimal stoichiometry: 1μM Hsc70, 5μM DnaJB1, and 0.1μM Apg2 (ns = p>0.05, all other values have p<0.05). **(C)** Heat map of data depicted in (A). The active region is outlined in red and represents the region with greater than 50% of the maximal activity when treated with DMSO. Color gradient represents luminescence. **(D)** Heat map of data depicted in (B). The active region is outlined in red and represents the region with greater than 50% of the maximal activity when treated with compound 18. Color gradient represents luminescence. **(E)** Difference heat map representing the change in luciferase disaggregation and reactivation between DMSO and compound 18 treated samples for each stoichiometric composition. Data were analyzed using unpaired t-test for each stoichiometric condition comparing DMSO treated (A) with compound 18 treated (B) (*p < 0.05).

The optimal concentration of Apg2 also changes with the concentration of DnaJB1. With 0.1μM DnaJB1 (Figure 5C), the optimal Apg2 concentration is 0.05μM. At higher DnaJB1 concentrations (≥0.5μM DnaJB1), the optimal Apg2 concentration is instead 0.1μM. These findings suggest that at low concentrations of DnaJB1, less Apg2 is required for robust disaggregase and reactivation activity. Likewise, at high DnaJB1 concentrations, more Apg2 is required for disaggregase and reactivation activity, and a greater concentration of Apg2 is tolerated before inhibiting the system. Thus, a balance between Hsp40 and Hsp110 supports optimal disaggregase activity by the Hsp70 system. Since Hsp40 promotes polypeptide capture by Hsp70 and Hsp110 promotes polypeptide release from Hsp70,^29^ this finding also suggests that an optimal proportion of Hsp70 must be in contact with the substrate for robust disaggregase activity.

115-7c mimics some aspects of Hsp40 activity.^49^ Thus, we hypothesized that the 115-7c analog, compound 18, would bolster Hsp40 activity within the disaggregase system. Indeed, compound 18 (25μM) greatly stimulated disaggregase and reactivation activity in specific regions of the landscape of chaperone stoichiometries (Figure 5B, D, E). These regions are readily visualized by a difference plot of the compound-treated versus vehicle-treated landscape (Figure 5E). The stoichiometry that is most robustly stimulated is 0.5μM DnaJB1 and 0.1μM Apg2 (Figure 5E). In fact, in the presence of 0.1μM Apg2, compound 18 stimulates activity in the presence of 0.1μM, 0.5μM, and 1μM DnaJB1 more than with 5μM DnaJB1 (Figure 5E). Since 1μM Hsc70, 5μM DnaJB1, and 0.1μM Apg2 is the optimal chaperone stoichiometry under vehicle-treated conditions (Figure 5A, C), these data indicate that chaperone stoichiometries with lower DnaJB1 concentrations are stimulated to a greater degree by compound 18. Thus, compound 18 increases the specific activity of DnaJB1.

This conclusion is further illustrated by charting the constellation of chaperone stoichiometries that achieve at least 50% of the maximal activity in the presence or absence of compound 18 (Figure 5C, D, red box). In the presence of compound 18, this region encompasses a greater area of the landscape and is shifted toward lower DnaJB1 concentrations (Figure 5C, D, red box). This shift suggests that compound 18 increases the effective DnaJB1 concentration yielding greater activity in regions where DnaJB1 concentrations are limiting. Furthermore, in the absence of DnaJB1, despite very minimal activity, we observe a slight but statistically significant increase in the activity of 1µM Hsc70 with 0.05µM or 0.1µM Apg2 consistent with compound 18 mimicking some aspects of Hsp40 function (Figure 5E).

Intriguingly, a portion of the landscape also emerges at a specific Apg2 concentration (0.01µM) where compound 18 inhibits activity (Figure 5E, red squares). At this Apg2 concentration (0.01µM), increasing DnaJB1 concentration above 0.5µM reduces activity (Figure 5A, C), and compound 18 exacerbates this effect (Figure 5B, D, E). Thus, here too, compound 18 seems to increase the effective DnaJB1 concentration. It is important to note that the region where compound 18 is inhibitory is only a small portion of the landscape (7/48 of the chaperone stoichiometries tested), and these differences are not statistically significant (Figure 5E). Conversely, compound 18 statistically significantly stimulates activity in ∼44% (21/48) of the chaperone stoichiometries assessed here (Figure 5E). Thus, compound 18 stimulates disaggregase activity in diverse positions within the landscape of chaperone stoichiometries. Collectively, our findings suggest a novel therapeutic approach to bolster the Hsp70-disaggregase machinery to combat aberrant protein aggregation in disease.

## Discussion

We have uncovered six dihydropyrimidines, 115-7c and five analogs – compounds 8, 16, 17, 18, and 19 – that significantly stimulate the human Hsp70-disaggregase system. Most notably, compound 18 stimulates the disaggregase activity of Hsc70, DnaJB1, and Apg2 up to ∼7-fold against disordered luciferase aggregates and ∼2-fold against αSyn PFFs. Importantly, compound 18 is not selective for Hsc70, DnaJB1, and Apg2 but stimulates other Hsp70, Hsp40, and Hsp110 groupings. Compound 18 most effectively stimulates chaperone sets containing combinations of Hsc70 or Hsp72, DnaJB1 or DnaJB4, and Apg2 or Hsp105. Strikingly, Hsc70 with DnaJA2 and either of the Hsp110s tested here disaggregates and reactivates luciferase, but compound 18 has no effect on these chaperone sets. This result suggests that compound 18 displays selectivity for class B Hsp40 proteins over class A Hsp40s, indicating that compound 18 may stimulate interactions between class B Hsp40s and Hsp70 that differ from interactions between class A Hsp40s and Hsp70. Interestingly, specific interactions between class B Hsp40 proteins and the C-terminal EEVD tetrapeptide tail of Hsp70 have been identified that are not involved in class A Hsp40 activity.^25^ We conclude that compound 18 stimulates the disaggregase activity of select Hsp70 and class B Hsp40 interacting pairs.

Class A and class B Hsp40 proteins can synergize to enhance the human Hsp70-disaggregase system.^15^ We establish that compound 18 stimulates Hsp70, Hsp40, and Hsp110 chaperone sets containing pairs of class A and class B Hsp40 proteins, as well as pairs of class B Hsp40 proteins, but not pairs of class A Hsp40 proteins. Our findings suggest that compound 18 does not greatly stimulate Hsp70, Hsp40, and Hsp110 chaperone sets that have minimal disaggregase activity against luciferase and only stimulates active sets. Thus, the intrinsic substrate selectivity of active chaperone sets is preserved upon pharmacological activation. A recent study revealed a vastly expanded interaction network for the Hsp70-Hsp40 machinery.^74^ An examination of whether each of these interactions synergistically enhance protein-disaggregase activity—as observed with DnaJA1, DnaJA2, DnaJB1, and DnaJB4—is an important future undertaking.

The identity of the chaperones within the Hsp70-disaggegase system as well as their relative stoichiometry dictate disaggregase activity. By exploring a matrix of 48 different chaperone stoichiometries, we found, unexpectedly, that the optimal chaperone concentrations for luciferase disaggregation and reactivation are 1μM Hsc70, 5μM DnaJB1, and 0.1μM Apg2. These results agree with prior studies that found disaggregase activity is optimal at substoichiometric ratios of Apg2 relative to Hsc70 and DnaJB1.^18^ However, others have reported optimal activity at an equimolar ratio of Hsp105 to Hsp72 and DnaJA1.^13^ This disparity may reflect the different Hsp70, Hsp40, and Hsp110 chaperones assessed, and highlights the importance of a more comprehensive study of the human Hsp70-disaggregase system. In both studies, Hsp40 concentrations were held constant at either half the Hsp70 concentration or half the total Hsp70 plus Hsp110 concentration.^13,18^ To our knowledge, our finding that excess Hsp40 is optimal for Hsp70, Hsp40, and Hsp110 disaggregase activity is unanticipated, and emphasizes the importance of high Hsp40 expression for optimal activity.

Interestingly, the disaggregation and reactivation activity does not correlate monotonically with DnaJB1 or Apg2 concentration. Rather, a balance between the two components dictates optimal disaggregase activity (Figure 6A). Indeed, we find very poor disaggregase activity in regions of the landscape in which there are high concentrations of DnaJB1 and low concentrations of Apg2, or high concentrations of Apg2 and low concentrations of DnaJB1. These findings suggest that while both Hsp40 and Hsp110 are required for disaggregase activity, excess of either component can be detrimental (Figure 6A). By comparing the landscape of chaperone stoichiometries in the presence or absence of compound 18, we establish that compound 18 operates to stimulate disaggregase activity by increasing the effective class B Hsp40 concentration. Remarkably, compound 18 significantly stimulates activity in ∼44% (21/48) of the chaperone stoichiometries assessed.

**Figure 6.**
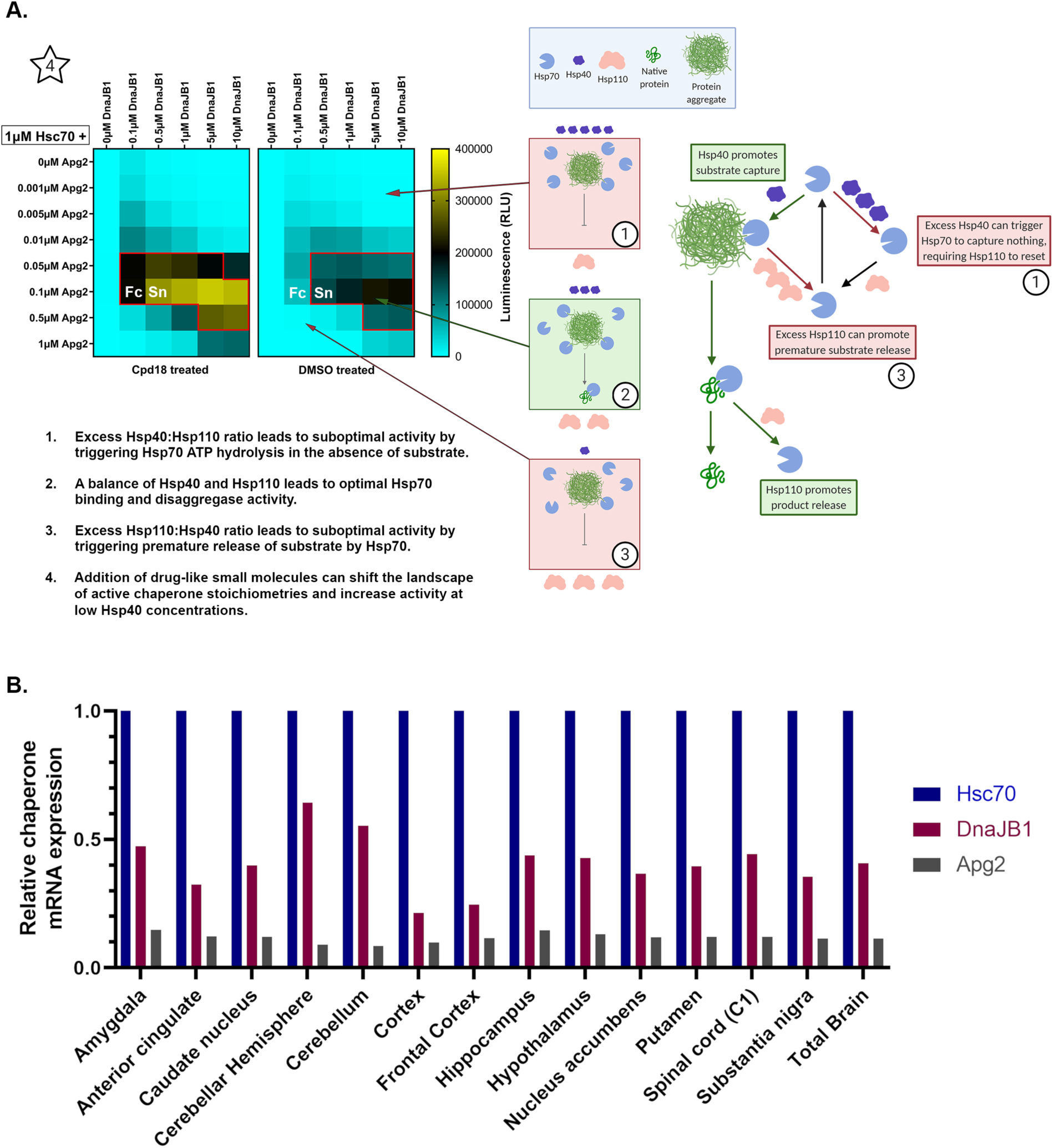
Compound 18 enhances the activity of the Hsp70-disaggregase system at putative chaperone stoichiometries found in the human brain. **(A)** Overview of compound 18 stimulation of the Hsp70-disaggregase system and proposed mechanistic interpretation of the stoichiometric landscape data. 1) Describes suboptimal scenarios with excess Hsp40, 2) describes the activity at optimal relative chaperone stoichiometry, and 3) describes suboptimal scenarios with excess Hsp110. The active regions are outlined in red and represent the regions with greater than 50% of the maximal activity for compound 18 or DMSO-treated conditions respectively. Fc and Sn (white) highlight the region corresponding to the chaperone ratios for the frontal cortex and substantia nigra, respectively, as determined in panel B. **(B)** Estimated relative mRNA expression of *Hsc70, DnaJB1, and Apg2* in various brain regions normalized to *Hsc70* values. Data sourced from the alternative splicing catalog of the transcriptome (ASCOT) database was used to estimate the relative expression of *Hsc70, DnaJB1, and Apg2* using publicly available RNA-seq datasets.^83^ Y-axis represents normalized mRNA expression estimates relative to *Hsc70*.

Hsp40 binds substrate and recruits Hsp70 and then stimulates ATP hydrolysis by Hsp70 causing a conformational shift in Hsp70 that results in substrate capture.^6,14,33,75–80^ Enhancing this step would yield more effective substrate capture and thus Hsp70 would be better primed for polypeptide extraction. When Hsp40 concentrations are in excess and Hsp110 levels are low, we observe a decline in disaggregase activity that may be explained by Hsp40 triggering Hsp70 to hydrolyze ATP in the absence of substrate, thus causing a conformation change in Hsp70 in the absence of substrate capture (Figure 6A). This futile step would require Hsp110 to reset Hsp70. In fact, we find that excess Hsp40 at low Hsp110 concentrations is detrimental to disaggregase activity.

Hsp110 is a NEF that induces ADP-ATP exchange in the Hsp70 NBD.^6,13,14,16–18,34,81^ Exchange of ADP for ATP releases the extracted polypeptide from Hsp70. Enhancing this step would improve the rate at which Hsp70 is reset and ready for another round of substrate capture and extraction. However, Hsp110 could also act on Hsp70 before polypeptide is extracted and thus promote premature substrate release, directly counteracting the effects of Hsp40. This possibility is supported by our results showing that at high Hsp110 concentrations there is an inhibition of disaggregase activity (Figure 6A).

What is the ratio of various Hsp70, Hsp40, and Hsp110 proteins in the brain? This question is important as imbalances in the concentrations of Hsp70 family members can foster tau accumulation.^82^ Moreover, drifts away from optimal chaperone stoichiometries could underlie selective vulnerability of some neuronal populations in neurodegenerative disease. To begin to address this question, we utilized the alternative splicing catalog of the transcriptome (ASCOT) database, which cross references tens of thousands of RNA-seq datasets to determine gene expression and splice frequency.^83^ The expression profile of *Hsc70*, *DnaJB1*, and *Apg2* mRNA across many regions of the CNS has an approximately 10:4:1 stoichiometric ratio, respectively (Figure 6B, Total Brain). If this stoichiometry is preserved at the protein level (but see caveats^84,85^), then this ratio is within the active region (i.e., having at least 50% of maximal activity) of the landscape of chaperone stoichiometries, yet is suboptimal (Figure 5C, 6A).

PD is characterized by the accumulation of Lewy bodies comprised of fibrillar αSyn within the dopaminergic neurons of the substantia nigra.^68^ Notably, the *Hsc70*:*DnaJB1*:*Apg2* mRNA ratio in the substantia nigra is approximately 10:3.5:1 (Figure 6B). Thus, DnaJB1 expression in the substantia nigra is slightly reduced compared to total brain but is near the ratio tested for αSyn disaggregation in Figure 3, indicating that compound 18 could improve the αSyn disaggregase activity of Hsc70, DnaJB1, and Apg2 in the substantia nigra. Moreover, we have shown that increasing DnaJB1 concentrations beyond that of Hsc70 can further increase the luciferase disaggregase and reactivation activity of Hsc70, DnaJB1, and Apg2. Hence, increasing DnaJB1 concentrations or activity in degenerating neurons could be a protective strategy for PD and other neurodegenerative disorders.^86–88^

Interestingly, the cortex and frontal cortex, which are affected in ALS/FTD, exhibit reduced levels of DnaJB1 with a *Hsc70*:*DnaJB1*:*Apg2* mRNA ratio is 10:2:1 (Figure 6B), which is on the border of the active region (Figure 5C red box, 6A white). This region of the landscape of chaperone stoichiometries is significantly bolstered by the addition of compound 18 (Figure 5D, 6A). Thus, compound 18 could similarly improve the disaggregase activity of Hsc70, DnaJB1, and Apg2 in the frontal cortex and potentially reduce aggregation of FUS, TAF15, or TDP-43 in ALS/FTD.^89^ Importantly, compound 18 significantly stimulates disaggregase activity in a large fraction of chaperone stoichiometries, including all of those measured for the various brain regions in the ASCOT dataset (Figure 6B).

Genetic studies suggest that altering specific chaperone components within the Hsp70-disaggregase system could be beneficial in models of neurodegenerative disease.^90^ For example, overexpression of Apg1 (HSPH3, an Hsp110) in a mutant SOD1^G85R^ mouse model of ALS showed improved survival.^91^ By contrast, Hsc70 overexpression in the SOD1^G85R^ mice did not extend survival, suggesting that Hsp110 may be the limiting chaperone factor in this model.^91^ Overexpression of Apg1 also reduced αSyn pathology in a transgenic mouse model expressing mutant αSyn^A53T^, a PD mouse model using injected αSyn PFFs, and in HEK293T cells overexpressing αSyn.^92^ In another study, knockdown of the *C. elegans* homologs of Hsp70 (hsp-1), Hsp40 (dnj-13), and Hsp110 (hsp-110) increased Htt-polyQ aggregation in this HD model.^21^ Knockdown of DnaJB1 or Apg2 in HD patient-derived neural progenitor cells also increased Htt-polyQ aggregation.^21^ Overall, these studies suggest that altering expression of components of the Hsp70-disaggregase system can mitigate protein aggregation pathology *in vivo*.

It is important to emphasize that manipulating the Hsp70-disaggregase system to confer neuroprotection is a delicate operation, and some alterations could be problematic. Indeed, overactivation of Hsp70 by specific Hsp40s can underlie disease.^93,94^ Small-molecule inhibitors of Hsp70, which would presumably reduce disaggregase activity, can prevent pathological tau accumulation.^95–99^ Moreover, the Hsp70-disaggregase system can promote protein aggregation and toxicity in *C. elegans.*^100^ Knockdown of *C. elegans* Hsp110 reduced luciferase disaggregation but also reduced αSyn and polyQ aggregation and toxicity.^100^ Here, it is suggested that the Hsp70-disaggregase system might promote prion-like propagation via enhanced fragmentation of amyloid fibrils.^100^ However, *in vitro* the Hsp70-disaggregase system preferentially liberates protein monomers from the ends of amyloid fibrils, which should minimize deleterious fibril fragmentation.^19,20,24,27,28,92^ Nonetheless, caution is warranted as it is important to ensure that a therapeutic regime of protein disaggregation is achieved such that protein solubilization is achieved rapidly without amplification of prion-like conformers that spread disease.^9^ The multicomponent nature and complexity of the Hsp70-disaggregase system makes this task challenging. For this reason, single-component disaggregases may prove to be more tractable therapeutically.^70,72,101–108^

Nevertheless, it may be advantageous to stimulate the activity of the Hsp70-disaggregase system in neurodegenerative disease in a controlled manner. A pharmacological approach has advantages in that treatment can more readily be administered transiently or intermittently to reduce unwanted effects.^9^ Compound 18 is a promising starting point towards developing pharmacological interventions that stimulate the human Hsp70-disaggregase system. We show here that several drug-like small molecules directly stimulate the disaggregase activity of the human Hsp70 system. However, some of the dihydropyrimidines used in this study are ineffective in H4 neuroglioma cell models.^46^ More specifically, 115-7c reduces αSyn aggregation in H4 neuroglioma cells,^55^ but compounds 16, 17, 18, 19, 20, 21, and 22 are ineffective.^46^ None of the dihydropyrimidines were toxic.^46^ Interestingly, compound 26 reduced αSyn aggregation in H4 neuroglioma cells,^46^ but did not enhance disaggregase activity in our studies (Figure 2A). One possibility is that compounds 16, 17, 18, 19, 20, 21, 22, and 26, which are esterified derivatives of 115-7c, could be differentially hydrolyzed by cellular carboxylesterases to yield 115-7c in the H4 model. It is also unknown if 115-7c or its derivatives pass the blood-brain-barrier, but they are predicted to not be able to cross (Table S1).^109^ Although compound 18 is ineffective in H4 neuroglioma cells, it provides proof-of-principle that human Hsp70-disaggregase activity can be pharmacologically stimulated. Ultimately, designing brain-penetrant, drug-like small molecules that mimic the effect of the active compounds discovered in this study represents a new strategy for the treatment of some classes of neurodegenerative diseases.

In summary, we establish that drug-like small molecules can be used to stimulate the disaggregase activity of the human Hsp70 system under a wide variety of chaperone stoichiometries. These findings suggest a therapeutic strategy to correct suboptimal proteostasis within aging neurons in neurodegenerative disease.^9^ Despite robust disaggregase activity *in vitro*, the human Hsp70-disaggregase system fails to counter protein aggregation in neurodegenerative disease. Our data raise the possibility that this deficit could be due to altered expression of Hsp70, Hsp40, or Hsp110 to yield suboptimal stoichiometries for disaggregation. Thus, it is critical to determine the expression levels of these chaperones in selectively vulnerable neurons in normal and disease conditions to establish if stoichiometries are altered. In addition, chaperone concentrations within the cell vary drastically in response to various stresses.^110,111^ Hsp70 expression is also altered as a function of aging and the same may occur for Hsp40 and Hsp110.^112,113^ One possibility is that as selectively vulnerable neurons age, the expression profiles of Hsp70, Hsp40, or Hsp110 chaperones drift to suboptimal stoichiometries. Thus, disaggregase activity would be reduced and allow formation, persistence, and propagation of aggregated conformers observed in patients with neurodegenerative diseases. However, compound 18 stimulates disaggregase activity at a wide spectrum of chaperone stoichiometries by increasing the effective activity of class B Hsp40s. Thus, finding a brain-penetrant analog of compound 18, which retains activity in neurons, could pharmacologically bolster the Hsp70-disaggregase to combat aberrant protein aggregation in disease. We suggest that further development of compound 18 will enable therapeutic strategies for several debilitating neurodegenerative disorders.

## STAR METHODS

Key resources table

| REAGENT or RESOURCE | SOURCE | IDENTIFIER |
| --- | --- | --- |
| **Antibodies** |  |  |
| alpha Synuclein Mouse Monoclonal Antibody (Syn 211) | ThermoFisher Scientific | Cat # AHB0261<br>RRID:AB_2536241 |
| IRDye 800CW Goat anti-Mouse IgG secondary antibody | LI-COR | Cat# 926–32210;<br>RRID:AB_621842 |
| **Bacterial and virus strains** |  |  |
| *E. coli* BL21 (DE3) RIL cells | Agilent | Cat # 230245 |
| **Chemicals, peptides, and recombinant proteins** |  |  |
| HisPur™ Ni-NTA Resin | ThermoFisher Scientific | Cat # 88223 |
| Resource Q column (6mL) | GE Healthcare | GE17-1179-01 |
| Resource S column (6mL) | GE Healthcare | GE17-1180-01 |
| Lysozyme | Sigma-Aldrich | Cat # L6876 |
| Firefly luciferase | Sigma-Aldrich | Cat # L-9506 |
| Creatine kinase | Roche | Cat # 10127566001 |
| Creatine phosphate | Roche | Cat # 10621722001 |
| cOmplete Mini, EDTA-free protease inhibitor | Roche | Cat # 11835170001 |
| ATP | Sigma-Aldrich | Cat # A3377 |
| His6-SUMO-Hsc70 | (Michalska et al., 2019)<sup>114</sup> | N/A |
| His6-SUMO-Hsp72 | This paper | N/A |
| His6-SUMO-DnaJA1 | (Michalska et al., 2019)<sup>114</sup> | N/A |
| His6-SUMO-DnaJA2 | This paper | N/A |
| His6-SUMO-DnaJB1 | (Michalska et al., 2019)<sup>114</sup> | N/A |
| His6-SUMO-DnaJB3 | This paper | N/A |
| His6-SUMO-DnaJB4 | This paper | N/A |
| His6-SUMO-Apg2 | This paper | N/A |
| His6-SUMO-Hsp105 | This paper | N/A |
| PARL<sup>Skd3</sup> | (Cupo et al., 2020)<sup>72</sup> | N/A |
| Hsp104<sup>K358D</sup> | (Mack et al., 2023)<sup>70</sup> | N/A |
| Brij™-35, 30% Solution | Thermo Scientific | Cat # 20150 |
| Alpha synuclein PFFs | (Luk et. al., 2012)<sup>68</sup> | N/A |
| Cpd-1 (JRB-473-66, MAL1-56B) | This paper | N/A |
| Cpd-2 (JRB-473-67, MAL1-56A) | This paper | N/A |
| Cpd-3 (JRB-473-68) | This paper | N/A |
| Cpd-4 (JRB-473-69, MAL2-101) | This paper | N/A |
| Cpd-5 (MLR633-018) | This paper | N/A |
| 115-7c (MAL1-271) | StressMarq | SKU: SIH-123-25MG |
| Cpd-7 (MAL1-55D) | Aurora Fine Chemicals | Cat # 153.627.403 |
| Cpd-8 (MAL2-06A) | (Werner et al., 2006)<sup>57</sup> | N/A |
| Cpd-9 (MAL3-101) | (Werner et al., 2006)<sup>57</sup> | N/A |
| Cpd-10 (DMT-022-20) | (Werner et al., 2006)<sup>57</sup> | N/A |
| Cpd-11 (DMT-022-22) | (Werner et al., 2006)<sup>57</sup> | N/A |
| Cpd-12 (DMT-022-86) | (Werner et al., 2006)<sup>57</sup> | N/A |
| Cpd-13 (DMT-031-10) | (Werner et al., 2006)<sup>57</sup> | N/A |
| Cpd-14 (DMT-031-12) | (Werner et al., 2006)<sup>57</sup> | N/A |
| Cpd-15 (MAL2-11B tetrazole) | (Huryn et al., 2011).<sup>58</sup> | N/A |
| Cpd-16 (AMT-628-27) | (Chiang et al., 2019)<sup>46</sup> | N/A |
| Cpd-17 (DWN-723-28) | (Chiang et al., 2019)<sup>46</sup> | N/A |
| Cpd-18 (DWN-723-35) | (Chiang et al., 2019)<sup>46</sup> | N/A |
| Cpd-19 (DWN-723-36) | (Chiang et al., 2019)<sup>46</sup> | N/A |
| Cpd-20 (DWN-723-38) | (Chiang et al., 2019)<sup>46</sup> | N/A |
| Cpd-21 (DWN-723-39) | (Chiang et al., 2019)<sup>46</sup> | N/A |
| Cpd-22 (DWN-723-40) | (Chiang et al., 2019)<sup>46</sup> | N/A |
| Cpd-23 (ML282-86) | (Ireland et al., 2014)<sup>59</sup> | N/A |
| Cpd-24 (TSM-592-54) | (Maskrey et al.,<br>2018)<sup>62</sup> | N/A |
| Cpd-25 (SHM-027-13) | This paper | N/A |
| Cpd-26 (DWN-723-23) | (Chiang et al., 2019)<sup>46</sup> | N/A |
| Cpd-27 (CBRC1039158) | Aurora Fine Chemicals | Cat # 174.670.865 |
| Cpd-28 (MAL1-47C) | Aurora Fine Chemicals | Cat # 153.735.968 |
| Cpd-29 (TSM-592-59) | (Maskrey et al.,<br>2018)<sup>62</sup> | N/A |
| Cpd-30 (ENAT5875208) | Enamine | Product ID<br>Z47327321 |
| Cpd-31 (DML-044-10) | This paper | N/A |
| Cpd-32 (ENAT5825922) | Enamine | Product ID<br>Z46740156 |
| **Critical commercial assays** |  |  |
| ATPase Activity Kit (Colorimetric) | Innova Biosciences | Cat # 601–0120 |
| Luciferase Assay Reagent | Promega | Cat # E1483 |
| **Deposited data** |  |  |
| Alternative splicing catalog of the transcriptome (AS-COT) | (Ling et al., 2020)<sup>83</sup> | N/A |
| **Recombinant DNA** |  |  |
| pE-SUMO Vector | LifeSensors | Cat # 1001A |
| Hsc70 in pE-SUMO | (Michalska et al., 2019)<sup>114</sup> | N/A |
| Hsp72 in pE-SUMO | This paper | N/A |
| DnaJA1 in pE-SUMO | (Michalska et al., 2019)<sup>114</sup> | N/A |
| DnaJA2 in pE-SUMO | This paper | N/A |
| DnaJB1 in pE-SUMO | (Michalska et al., 2019)<sup>114</sup> | N/A |
| DnaJB3 in pE-SUMO | This paper | N/A |
| DnaJB4 in pE-SUMO | This paper | N/A |
| Apg2 in pE-SUMO | This paper | N/A |
| Hsp105 in pE-SUMO | This paper | N/A |
| PARL Skd3 in pMAL C2 with TEV site | (Cupo et al., 2020)<sup>72</sup> | N/A |
| pNOTAG-Hsp104<sup>K358D</sup> | (Mack et al., 2023)<sup>70</sup> | N/A |
| pFGET19 Ulp1 | Addgene | Plasmid #64697 |
| **Software and algorithms** |  |  |
| Prism 7/8/9 | GraphPad | N/A |
| ImageJ | (Schindelin et al., 2012)<sup>115</sup> | N/A |

### RESOURCE AVAILABILITY

#### Lead contact

Further information and requests for resources and reagents should be directed to and will be fulfilled by the lead contact, James Shorter.

#### Materials availability

Plasmids or compounds newly generated in this study will be made readily available to the scientific community. We will honor requests in a timely fashion. Material transfers will be made with no more restrictive terms than in the Simple Letter Agreement or the Uniform Biological Materials Transfer Agreement and without reach through requirements.

#### Data and code availability

Any additional information required to reanalyze the data reported in this paper is available from the lead contact upon request.

### EXPERIMENTAL MODEL AND SUBJECT DETAILS

*E. coli* BL21 (DE3) RIL cells from Agilent (Cat# 230245) were used for protein purification.

### METHODS DETAILS

#### Protein expression and purification

*Hsp72, Hsc70, DnaJA1, DnaJA2, DnaJB1, DnaJB3, DnaJB4, Hsp105, and Apg2* cDNA were obtained from Addgene or kindly gifted to us by Mikko Taipale from the University of Toronto. Proteins were expressed as N-terminally His_6_-SUMO tagged fusion proteins from pE-SUMOpro plasmid (Life Sensors) in BL21 DE3 RIL *E. coli*. Hsp110 proteins were cloned with a two-glycine linker between His_6_-SUMO and Hsp110 as previously described.^18,114^ Transformed bacteria were grown in Luria broth with 25µg/mL chloramphenicol and 100µg/mL ampicillin at 37°C with 250rpm shaking. At an OD_600_ of 0.6, protein expression was induced with 1mM IPTG for 16 hours at 15°C with 250rpm shaking. Cells were harvested and lysed in lysis buffer (50mM HEPES pH 7.5, 750mM KCl, 5mM MgCl_2_, 10% glycerol, 20mM imidazole, 2mM β-mercaptoethanol, 5µM pepstatin A, and Roche cOmplete mini EDTA-free protease inhibitor) by treatment with 66μg/mL lysozyme followed by sonication. Lysates were clarified by ultracentrifugation at 30597xg for 20 min. Then cleared lysates were incubated with Thermo HisPur Ni-NTA resin for 90min at 4°C. The resin was then washed with 10 bead volumes of wash buffer (50mM HEPES pH 7.5, 750mM KCl, 5mM MgCl_2_, 10% glycerol, 20mM imidazole, 2mM β-mercaptoethanol, and 1mM ATP) and eluted with 2 bead volumes of elution buffer (50mM HEPES pH 7.5, 750mM KCl, 5mM MgCl_2_, 10% glycerol, 300mM imidazole, 2mM β-mercaptoethanol, and 1mM ATP). The protein was then cleaved by a 100:1 molar ratio of target protein to His-tagged Ulp1 overnight at 4°C concurrently with dialysis in wash buffer. The His-SUMO tag and His-Ulp1 were then removed by incubating with Ni-NTA resin for 90min at 4°C and collecting the supernatant.

The proteins were then further purified by ion exchange using either 6mL Resource Q (anion exchange) or 6mL Resource S (cation exchange) resin depending on the charge of the protein. For anionic Hsc70, Hsp72, DnaJA1, DnaJA2, DnaJB3, Apg2, and Hsp105, the protein was diluted 10-fold with Q0 buffer (20mM Tris pH 8.0, 0.5mM EDTA, 5mM MgCl_2_, 10% glycerol, 2mM β-mercaptoethanol, and 1mM ATP), and loaded onto the Resource Q column at 1mL/min. The column was then washed with 5 column volumes of Q50 buffer (20mM Tris pH 8.0, 0.5mM EDTA, 5mM MgCl_2_, 10% glycerol, 2mM β-mercaptoethanol, and 50mM NaCl), followed by a 0% to 50% buffer elution gradient of Q1000 buffer (20mM Tris pH 8.0, 0.5mM EDTA, 5mM MgCl_2_, 10% glycerol, 2mM β-mercaptoethanol, and 1000mM NaCl) over 10 column volumes. Fractions containing the target protein were pooled, buffer exchanged into storage buffer (40mM HEPES pH 7.4, 150mM KCl, 20mM MgCl_2_, 10% glycerol, 1mM DTT), and snap frozen in liquid nitrogen. For cationic DnaJB1 and DnaJB4, the proteins were treated the same, except using the Resource S column and S0 (20mM MES pH 6.0, 0.5mM EDTA, 5mM MgCl_2_, 10% glycerol, 2mM β-mercaptoethanol, and 1mM ATP), S50 (20mM MES pH 6.0, 0.5mM EDTA, 5mM MgCl_2_, 10% glycerol, 2mM β-mercaptoethanol, and 50mM NaCl), and S1000 (20mM MES pH 6.0, 0.5mM EDTA, 5mM MgCl_2_, 10% glycerol, 2mM β-mercaptoethanol, and 1000mM NaCl) buffers.

pFGET19 Ulp1 was obtained from Addgene. Ulp1 was expressed as an N-terminally His_6_-tagged fusion protein in BL21 DE3 RIL *E. coli*. Transformed bacteria were grown in Luria broth with 25µg/mL chloramphenicol and 50µg/mL kanamycin at 37°C with 250rpm shaking. At an OD_600_ of 0.6, protein expression was induced with 1mM IPTG for 16 hours at 15°C with 250rpm shaking. Cells were harvested and lysed in lysis buffer (50mM phosphate buffer pH 8.0, 300mM NaCl, 20mM imidazole, 2mM β-mercaptoethanol, 5 µM pepstatin A, and Roche cOmplete mini EDTA-free protease inhibitor) by treatment with 66μg/mL lysozyme followed by sonication. Lysates were clarified by ultracentrifugation at 30597xg for 20 min. Then cleared lysates were incubated with Thermo HisPur Ni-NTA resin for 90min at 4°C. The resin was then washed with 10 bead volumes of wash buffer (50mM phosphate buffer pH 8.0, 300mM NaCl, 20mM imidazole, 2mM β-mercaptoethanol) and eluted with 3 bead volumes of elution buffer (50mM phosphate buffer pH 8.0, 300mM NaCl, 250mM imidazole, 2mM β-mercaptoethanol). An equal volume of glycerol was added to the eluant and stored for short-term use at -20°C or -80°C for long-term storage.

Hsp104^K358D^ was purified as described.^70^ _PARL_Skd3 and TEV protease were purified as described.^6,116^

### Small molecules

115-7c was purchased from StressMarq. Reference compounds were synthesized as previously described.^46,57–59,62,117^ Compounds 1, 2, 3, 4, 5, 25 and 31 were prepared analogously; the experimental details and characterization data are listed below. Compounds 7 (Aurora Fine Chemicals), 27 (Aurora Fine Chemicals), 28 (Aurora Fine Chemicals), 30 (Enamine), and 32 (Enamine) are commercially available from the indicated as well as other suppliers. All samples passed QC with LCMS purities >95% before testing.

4-(5-((Benzyloxy)carbonyl)-4-(2-chlorophenyl)-6-methyl-2-oxo-3,4-dihydropyrimidin-1(2*H*)-yl)butanoic acid (compound 1). A solution of 4-ureidobutanoic acid (0.125g, 0.855mmol, 1 eq), 2-chlorobenzaldehyde (0.180g, 1.28mmol, 1.5 eq), and THF (2mL) was treated with benzyl acetoacetate (0.222mL, 1.28mmol, 1.5 eq) and conc. HCl (2 drops), stirred overnight at room temperature, and concentrated in vacuo. The residue was washed with hexanes, and dried to yield compound 1 (0.310g, 0.700 mmol, 82%) as a crystalline solid: Mp 181.0-181.3 °C; ATM-IR (neat) 1704, 1629, 1215, 1163, 1094, 748 cm-1; ^1^H NMR (300 MHz, DMSO-d6) ο 12.12 (bs, 1 H), 7.90 (d, *J* = 3.3 Hz, 1 H), 7.42-7.39 (m, 1 H), 7.27-7.22 (m, 7 H), 7.03-7.00 (m, 2 H), 5.64 (d, *J* = 3.3 Hz, 1 H), 5.00, 4.99 (AB, *J* = 12.9 Hz, 2 H), 3.88-3.80 (m, 1 H), 3.63-3.55 (m, 1 H), 2.59 (s, 3 H), 2.23 (t, *J* = 6.9 Hz, 2 H), 1.81-1.66 (m, 2 H); ^13^C NMR (125 MHz, DMSO-d6) ο 173.9, 165.0, 151.8, 151.5, 140.5, 136.3, 132.0, 129.7, 129.3, 128.3, 128.2, 127.7, 127.6, 127.3, 100.9, 65.0, 50.2, 41.3, 30.8, 24.6, 15.6; HRMS (ESI) m/z calcd for C_23_H_24_N_2_O_5_Cl ([M+1]^+^) 443.1368, found 443.1366.

4-(5-((Benzyloxy)carbonyl)-4-(4-chlorophenyl)-6-methyl-2-oxo-3,4-dihydropyrimidin-1(2*H*)-yl)butanoic acid (compound 2). A solution of 4-ureidobutanoic acid (0.125g, 0.855 mmol, 1 eq), 4-chlorobenzaldehyde (0.222mL, 0.180g, 1.28mmol, 1.5 eq), and THF (1mL) was treated with benzyl acetoacetate (0.222mL, 1.28mmol, 1.5 eq) and conc. HCl (2 drops), stirred overnight at room temperature, and concentrated in vacuo. The residue was washed with hexanes, and dried to yield compound 2 (0.313g, 0.707mmol, 83%) as a crystalline solid: Mp 195.1-197.1 °C; ATM-IR (neat) 1704, 1629, 1420, 1232, 1163, 1094, 749 cm-1; ^1^H NMR (300 MHz, DMSO-d6) 8 12.11 (bs, 1 H) 7.97 (d, *J* = 3.6 Hz, 1 H), 7.33-7.24 (m, 7 H), 7.18-7.13 (m, 2 H), 5.15 (d, *J* = 3.6 Hz, 1 H), 5.07, 5.02 (AB, *J* = 12.6 Hz, 2 H), 3.83-3.75 (m, 1 H), 3.57-3.44 (m, 1 H), 2.59 (s, 3 H), 2.13 (t, *J* = 6.9 Hz, 2 H), 1.74-1.53 (m, 2 H); ^13^C NMR (125 MHz, DMSO-d6) δ 173.9, 165.2, 152.4, 150.8, 142.7, 136.3, 132.0, 128.5, 128.3, 128.1, 127.9, 127.7, 102.2, 65.3, 51.9, 41.2, 30.6, 24.6, 15.7; HRMS (ESI) m/z calcd for C_23_H_24_N_2_O_5_Cl ([M+1]^+^) 443.1368, found 443.1366.

4-(5-((Benzyloxy)carbonyl)-4-(4-fluorophenyl)-6-methyl-2-oxo-3,4-dihydropyrimidin-1(2*H*)-yl)butanoic acid (compound 3). A solution of 4-ureidobutanoic acid (0.125 g, 0.855 mmol, 1 eq), 4-fluorobenzaldehyde (0.159g, 1.28mmol, 1.5 eq), and THF (1mL) was treated with benzyl acetoacetate (0.222mL, 1.28mmol, 1.5 eq) and conc. HCl (2 drops), stirred overnight at room temperature, and concentrated in vacuo. The residue was washed with hexanes, and dried to give compound 3 (0.308g, 0.722mmol, 84%) as a crystalline solid: Mp 200.2-202.2 °C; ATM-IR (neat) 1704, 1629, 1420, 1232, 1215, 1163, 839 cm^-1^; ^1^H NMR (300 MHz, DMSO-d6) δ 12.09 (bs, 1 H) 7.98 (d, *J* = 3.9 Hz, 1 H), 7.31-7.10 (m, 9 H), 5.18 (d, *J* = 3.9 Hz, 1 H), 5.09, 5.05 (AB, *J* = 12.6 Hz, 2 H), 3.86-3.78 (m, 1 H), 3.60-3.49 (m, 1 H), 2.59 (s, 3 H), 2.13 (t, *J* = 7.2 Hz, 2 H), 1.80-1.50 (m, 2 H); ^13^C NMR (125 MHz, DMSO-d6) δ 173.8, 165.2, 161.3 (d, *J* =241.3 Hz), 152.4, 150.6, 140.0 (d, *J* = 2.5 Hz), 136.3, 128.3, 128.1, 128.0, 127.8, 127.6, 115.1 (d, *J* = 21.3 Hz), 102.5, 65.1, 51.7, 41.1, 30.5, 24.5, 15.6; HRMS (ESI) *m/z* calcd for C_23_H_24_N_2_O_5_F ([M+1]^+^) 427.1664, found 427.1662.

4-(5-((Benzyloxy)carbonyl)-4-(4-(trifluoromethyl)phenyl)-6-methyl-2-oxo-3,4-dihydropyrimidin-1(2*H*)-yl)butanoic acid (compound 4). A solution of 4-ureidobutanoic acid (0.125g, 0.855mmol, 1 eq), 4-(trifluoromethyl)benzaldehyde (0.175mL, 0.223g, 1.28mmol, 1.5 eq), and THF (1mL) was treated with benzyl acetoacetate (0.222mL, 1.28mmol, 1.5 eq) and conc. HCl (2 drops), stirred overnight at room temperature, and concentrated in vacuo. The residue was washed with hexanes, and dried to give compound 4 (0.317g, 0.665mmol, 78%) as a crystalline solid: Mp 196.6-198.6 °C; ATM-IR (neat) 1704, 1632, 1420, 1300, 1232, 1109 cm^-1^; ^1^H NMR (300 MHz, DMSO-d6) δ 12.08 (bs, 1 H) 8.06 (d, *J* = 3.9 Hz, 1 H), 7.65 (d, *J* = 8.1 Hz, 2 H), 7.39 (d, *J* = 8.1 Hz, 2 H), 7.27-7.14 (m, 3 H), 7.14-7.12 (m, 2 H), 5.26 (d, *J* = 3.3 Hz, 1 H), 5.10, 5.03 (AB, *J* = 12.6 Hz, 2 H), 3.86-3.78 (m, 1 H), 3.58-3.50 (m, 1 H), 2.54 (s, 3 H), 2.12 (t, *J* = 7.2 Hz, 2 H), 1.80-1.55 (m, 2 H); ^13^C NMR (125 MHz, DMSO-d6) δ 173.9, 165.1, 152.3, 151.2, 148.3, 136.3, 128.5, 128.3, 128.1 (q, *J* = 31.3 Hz), 127.8, 127.7, 127.4, 127.0, 125.5 (q, *J* = 3.8 Hz), 124.2 (q, *J* = 270.0 Hz), 101.8, 65.2, 52.2, 41.3, 30.6, 24.6, 15.7; HRMS (ESI) *m/z* calcd for C_24_H_24_N_2_O_5_F_3_ ([M+1]^+^) 477.1632, found 477.1630.

4-(5-((Benzyloxy)carbonyl)-4-(2,4-dichlorophenyl)-6-methyl-2-oxo-3,4-dihydropyrimidin-1(2*H*)-yl)butanoic acid (compound 5). A solution of 3-ureidobenzoic acid (0.125g, 0.694mmol, 1 eq), 2,4-dichlorobenzaldehyde (0.123g, 0.694mmol, 1 eq), and THF (2mL) was treated with benzyl acetoacetate (0.137g, 0.694mmol, 1 eq) and conc. HCl (2 drops), stirred for 24h at room temperature, and concentrated in vacuo. The residue was purified by chromatography on SiO2 (EtOAc:hexanes, 2:1 to 3:1) to give crude product that was washed with hexanes and dried in vacuo to give compound 5 (0.59g, 0.311mmol, 45%) as a crystalline solid: Mp 236.0-236.8 °C (dec.); ATM-IR (neat) 1686, 1439, 1286, 1216, 1147, 1071, 751, 695 cm^-1^; ^1^H NMR (500 MHz, DMSO-d6/D_2_O) δ 7.95 (d, *J* = 8.0 Hz, 1 H), 7.71 (bs, 1 H), 7.57 (t, *J* = 7.7 Hz, 1 H), 7.49-7.45 (m, 2 H), 7.43 (d, *J* = 8.0 Hz, 1 H), 7.35 (d, *J* = 8.0 Hz, 1 H), 7.19-7.15 (m, 3 H), 6.91 (d, *J* = 7.0 Hz, 1 H), 5.73 (s, 1 H), 5.03, 4.87 (AB, *J* = 12.8 Hz, 2 H), 3.88-3.80 (m, 1 H), 2.50 (s, 3 H); ^13^C NMR (100 MHz, DMSO-d6) δ 166.7, 164.7, 150.9, 150.6, 139.7, 137.9, 136.1, 133.1, 133.0, 131.8, 130.4, 129.3, 129.1, 128.2, 128.1, 127.8, 127.5, 100.9, 65.2, 50.6, 18.3; HRMS (ESI) *m/z* calcd for C_26_H_21_N_2_O_5_Cl_2_ ([M+1]^+^) 511.0822, found 511.0825.

Benzyl 1-(4-((1-(butylamino)-1-oxopropan-2-yl)(2-morpholinoethyl)amino)-4-oxobutyl)-6-methyl-4-(4-nitrophenyl)-2-oxo-1,2,3,4-tetrahydropyrimidine-5-carboxylate (compound 25). A 5-mL microwave vial equipped with a stir bar was charged with 4-(5-((benzyloxy)carbonyl)-6-methyl-4-(4-nitrophenyl)-2-oxo-3,4-dihydropyrimidin-1(2*H*)-yl)butanoic acid^61^ (0.20g, 0.44mmol), MeOH (4.4mL), and 2-morpholinoethan-1-amine (0.060mL, 0.49mmol). The reaction mixture was stirred at 0 °C for 5 min and treated with *n*-butyl isocyanide (0.046mL, 0.44mmol) and acetaldehyde (0.25mL, 4.4mmol). The vial was capped with a microwave cap and was heated in a microwave reactor at 70°C for 60min. After cooling to room temperature, the brown solution was concentrated in vacuo and the crude residue was redissolved in CH_2_Cl_2_ and extracted with 10% NaOH (1x). The aqueous phase was extracted with CH_2_Cl_2_ (2x) and the combined organic layers were dried (Na_2_SO_4_) and concentrated *in vacuo* to give a crude residue that was purified by chromatography on SiO_2_ (CH_2_Cl_2_:MeOH, 100:0 to 90:10) to afford compound 25 (91.7mg, 28%) as a yellow, oily mixture of rotamers: IR 3272, 2958, 1685, 1522, 1388, 1158 cm^-1^; Major rotamer: ^1^H NMR (300 MHz, CDCl_3_) δ 8.04 (d, *J* = 8.1 Hz, 2 H), 7.33-7.27 (m, 5 H), 7.15-7.13 (m, 2 H), 6.68 (bs, 1 H), 6.42 (s, 1 H), 5.44 (s, 1 H), 5.12 (d, *J* = 12.0 Hz, 1 H), 5.01 (d, 1 H, *J* = 12.0 Hz), 4.09 (bs, 1 H), 3.88-3.56 (m, 6 H), 3.38-3.30 (m, 2 H), 3.25-3.13 (m, 2 H), 2.60 (s, 3 H), 2.53-2.43 (m, 7 H), 2.19 (bs, 1 H), 1.95-1.78 (m, 2 H), 1.46-1.19 (m, 11 H), 0.93-0.87 (m, 3 H). Characteristic signals of the minor rotamer: ^1^H NMR (300 MHz, CDCl_3_) δ 8.30 (bs, 1 H), 6.56 (bs, 1 H), 4.77-4.68 (m, 1 H), 2.57 (s, 3 H); HRMS (ESI) *m/z* calcd for C_36_H_49_N_6_O_8_ ([M+1]^+^) 693.3606, found 693.3583.

Benzyl 1-(4-((1-(butylamino)-1-oxopropan-2-yl)(2-(dimethylamino)ethyl)amino)-4-oxobutyl)-4-(4-chlorophenyl)-6-methyl-2-oxo-1,2,3,4-tetrahydropyrimidine-5-carboxylate (compound 31). According to the protocol used for compound 25, 4-(5-((benzyloxy)carbonyl)-4-(4-chlorophenyl)-6-methyl-2-oxo-3,4-dihydropyrimidin-1(2*H*)-yl)butanoic acid (0.254g, 0.573mmol), *N,N*-dimethylethylenediamine (0.069mL, 0.637mmol), *n*-butylisocyanide (0.067mL, 0.637mmol) and acetaldehyde (0.360mL, 6.37mmol) in MeOH (2mL) afforded compound 31 (137mg, 37%) as a yellow, oily mixture of rotamers: Major rotamer: ^1^H NMR (300 MHz, CDCl_3_) δ 9.21 (bs, 1 H), 7.31-7.07 (m, 9 H), 5.51 (bs, 1 H), 5.32 (bs, 1 H), 5.09, 5.02 (AB, *J* = 12.3 Hz, 2 H), 4.05 (bs, 1 H), 3.88-3.56 (m, 2 H), 3.41-3.14 (m, 4 H), 2.58 (s, 3 H), 2.49-2.36 (m, 2 H), 2.27 (s, 4 H), 2.22 (s, 2 H), 2.00-1.86 (m, 2 H), 1.48-1.25 (m, 9 H), 0.89 (t, *J* = 7.2 Hz, 3 H). Characteristic signals of the minor rotamer: ^1^H NMR (300 MHz, CDCl_3_) δ 6.98 (bs, 1 H), 5.48 (bs, 1 H), 4.65-4.59 (m, 1 H), 2.59 (s, 3 H), 2.20 (s, 2 H), 0.88 (t, *J* = 7.2 Hz, 3 H).

### Luciferase disaggregation and reactivation assays

Luciferase aggregates were generated by incubating 6mg/mL of recombinant firefly luciferase (Sigma) in luciferase refolding buffer (LRB: 25mM HEPES-KOH pH 7.4, 150mM potassium acetate, 10mM magnesium acetate, 10mM DTT) with 6M urea at 30°C for 30min. Denatured luciferase was then diluted 100-fold on ice into LRB (without urea), snap frozen, and stored at - 80°C until use.

Luciferase disaggregation and reactivation assays were setup as previously described.^107,118^ The chaperones and concentrations are indicated in each figure legend. Chaperones were incubated with 100nM luciferase aggregates (monomeric concentration) with an ATP regeneration system (ARS: 10mM creatine phosphate, 5mM ATP, 20μg/mL creatine kinase) in LRB. For experiments with small molecules, 0.001% Brij35 (w/v) and 1% final DMSO (v/v) were included in the LRB, and the concentration of the compound used is indicated in the figure legend. Samples were incubated at 25°C for 90 min and then mixed with luciferase assay reagent (Promega) and luminescence was measured in a Safire Tecan.

In luciferase disaggregation assays without compounds (i.e., no DMSO) chaperone concentrations are 1μM Hsp70, 0.5μM Hsp40, and 0.1μM Hsp110 (unless otherwise stated) and Hsc70, DnaJB1, and Apg2 recover approximately 15-30% of native luciferase. In assays with compounds (i.e., with 0.001% Brij35 (w/v) and 1% final DMSO (v/v) chaperone concentrations are 0.4μM Hsp70, 0.2μM Hsp40, and 0.04μM Hsp110 (unless otherwise stated) and Hsc70, DnaJB1, and Apg2 recover approximately 5% of native luciferase in DMSO control and 10-20% when treated with compound 18.

### ATPase assay

The steady state ATPase assay was performed as previously described.^107,118^ For Figure 2E, 0.4µM Hsc70, 0.2µM DnaJB1, and 0.04µM Apg2 were added to LRB with 0.001% Brij35 and either 25μM of the indicated compound or DMSO control (final 1% DMSO (v/v) in all reactions). For Figure S2, the chaperones and concentrations are indicated in the figure legend. The buffer used for ATPase assays in Figure S2 was 100mM Tris HCl pH 7.4, 20mM KCl, 6mM MgC_l2_, and 5mM DTT.

Added to the samples were 1mM ATP to start the reactions. At 0 min and 60 min, samples were taken and mixed with Pi Lock Gold mix from a Colorimetric ATPase Activity Kit (Innova Biosciences). After 2 min, stabilizer from the kit was added to the reactions. The reactions were then incubated on ice for 30 min before being read in a Safire Tecan for absorbance at 650nm.

### αSyn disaggregation assay

The αSyn disaggregation assay was performed as previously described.^72^ Briefly, 1μM Hsc70, 0.5μM DnaJB1, and 0.1μM Apg2 were added to LRB (25mM HEPES-KOH pH 7.4, 150mM potassium acetate, 10mM magnesium acetate, 10mM DTT) with 0.001% Brij35 and ARS (20mM creatine phosphate, 10mM ATP, 40μg/mL creatine kinase). Added to the samples were 0.5μM αSyn preformed fibrils kindly gifted to us from Kelvin Luk (University of Pennsylvania) and either the indicated concentration of compound 18 or DMSO control (final 1% DMSO (v/v) in all reactions).^68^ Samples were incubated at 37°C while shaking at 300rpm for 90min. Supernatant and pellet were generated by centrifugation at 20,000g for 20 min at 4°C. Pellets were resuspended in MSB (50mM Tris-HCl, pH 8.0, 8M Urea, 150mM NaCl). 10% of the total reaction, supernatant, or resuspended pellet were loaded onto nitrocellulose membrane using a 96-well vacuum manifold. Dot blots were then blocked, developed with mouse anti-SYN211 (Invitrogen) as the primary antibody and goat anti mouse as the secondary antibody, and imaged using an Odyssey Li-COR system. Images were analyzed using FIJI by measuring the integrated density of each dot. Soluble αSyn in the supernatant fraction was normalized by dividing by the total loaded αSyn for each corresponding condition and then plotted in GraphPad Prism.

### ASCOT database

Gene expression dataset for human tissues (GTEx) was downloaded as a .csv file from https://snaptron.cs.jhu.edu/data/ascot/. Data were sorted for brain tissues and members of the *Hsp70, Hsp40 (DnaJA* and *DnaJB),* and *Hsp110* chaperone families. Pseudogenes and *DnaJC* members were excluded from our gene expression analysis. Total Brain values are the average across all the brain tissues listed. Normalized area under the curve is an estimate of gene expression and is described here: http://ascot.cs.jhu.edu/naucpsi.html.

### Statistical methods

All statistical analyses were performed using GraphPad Prism version 7, 8, or 9. GraphPad Prism was used to calculate the % of maximal effect of compound 18 in Figure 3A using non-linear dose-response curve fitting. GraphPad Prism was used to analyze data using the multiple unpaired t-test and the one-way ANOVA followed by Dunnett’s or Tukey’s multiple comparisons test as indicated in figure legends.

## Acknowledgements

We thank Kelvin Luk, Mikko Taipale, and Amber Buhagiar for generously providing key reagents, and Danelle Lehsten for the preparation of compound 31. We thank JiaBei Lin and Edward Barbieri for feedback on the manuscript. Our work was supported by the Blavatnik Family Foundation Fellowship (EC & RRC), an ALS Association Award (JS), the Office of the Assistant Secretary of Defense for Health Affairs through the Amyotrophic Lateral Sclerosis Research Program (W81XWH-17-1-0237) (JS), NSF graduate research fellowship (DGE-1321851) (KLM), and NIH grants T32GM008076 (EC), T32GM008275 (RRC), F31AG060672 (RRC), K12GM081259 (ELG), P50GM067082 (PW), R35GM131732 (JLB), R21AG065854 (JS), R21AG061784 (JS), and R01GM099836 (JS).

---

**Figure S1.**
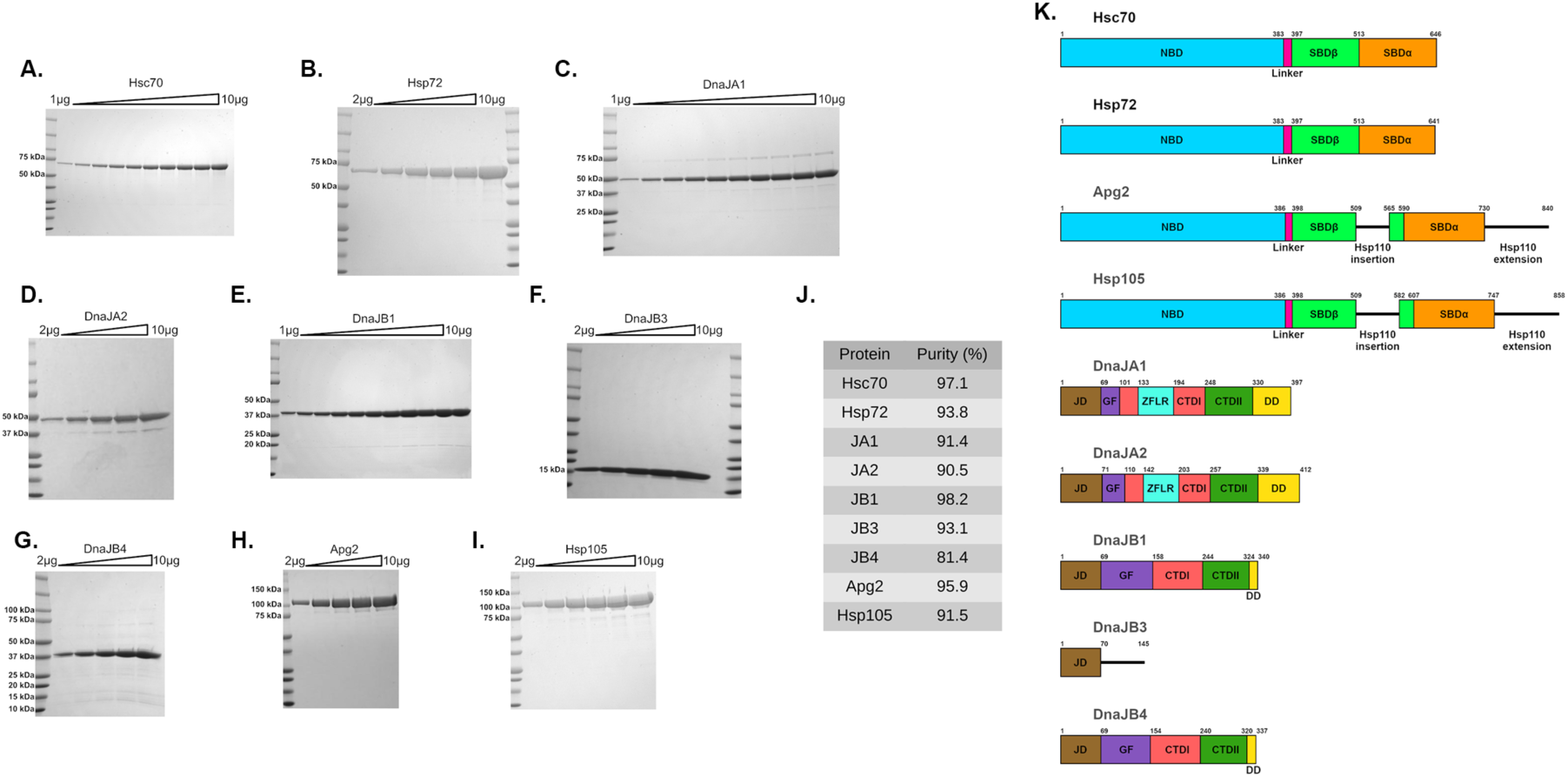
Purification and domain architecture of chaperones used in this study. **(A-I)** SDS-PAGE gels for all the chaperones purified. Samples were loaded from left to right with 1-10μg of protein in 1μg increments (A, C, and E). Alternatively, samples were loaded from left to right with 2-10μg of protein in 2μg increments (B, D, F, G, H, and I). Gels were stained with Coomassie Blue and imaged. **(J)** Purity was measured using ImageJ densitometry. Intensity of the band of interest was divided by the sum of the intensities for all the bands to calculate purity (%). **(K)** Domain maps of all the chaperones purified here. Key: nucleotide-binding domain (NBD), substrate-binding domain (SBD), J-domain (JD), G/F-rich region (GF), zinc-finger-like region (ZFLR), C-terminal domain (CTD), dimerization domain (DD). Domain start and end residues determined using clustal omega multiple sequence alignment and previously reported domain maps.^25,119–121^ Related to Figure 1.

**Figure S2.**
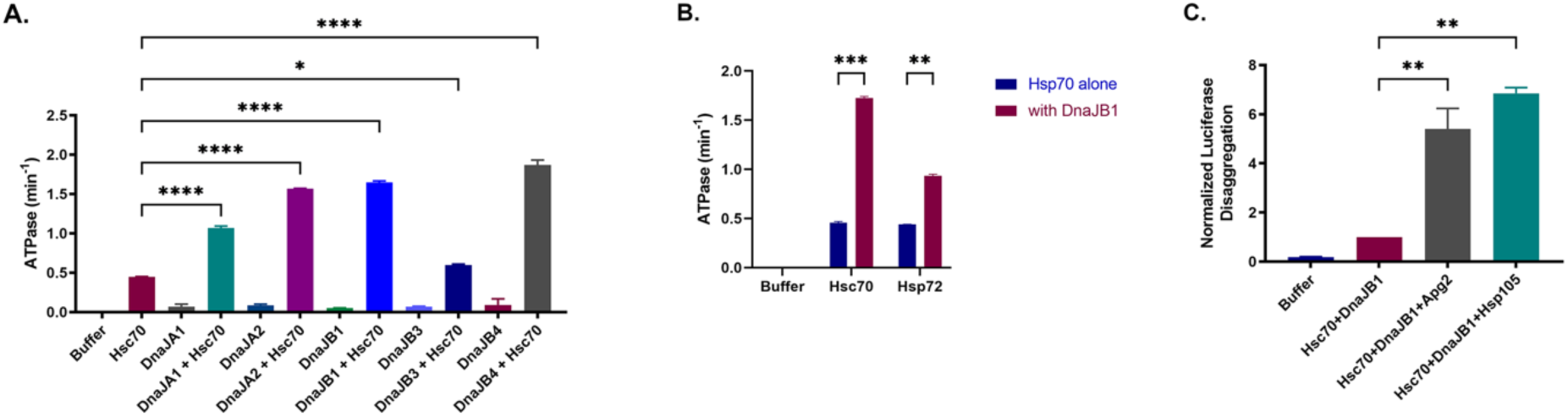
Purified chaperones are functional. **(A)** ATPase activity of each Hsp40 purified with or without Hsc70. 0.5μM of the indicated Hsp40 plus or minus 1μM Hsc70 were incubated with 1mM ATP. Values represent means ± SEM (n=2). Data were analyzed using one-way ANOVA followed by Dunnett’s MCT compared to Hsc70 alone (*p < 0.05, ****p < 0.001). **(B)** ATPase activity of each Hsp70 with and without DnaJB1. 1μM of the indicated Hsp70 plus or minus 0.5μM DnaJB1 were incubated with 1mM ATP. Values represent means ± SEM (n=2). Data were analyzed using multiple unpaired t-test between an Hsp70 protein with and without DnaJB1 (**p < 0.01, ***p < 0.005). **(C)** Luciferase disaggregase and reactivation activity of each Hsp110 purified with Hsc70 and DnaJB1. 1μM Hsc70, 0.5μM DnaJB1, and 0.1μM of the indicated Hsp110 were combined with 100nM luciferase aggregates (monomeric concentration) and an ATP-regenerating system. Values are normalized to Hsc70 and DnaJB1 without Hsp110 and are means ± SEM (n=2). Data were analyzed using one-way ANOVA followed by Dunnett’s MCT compared to Hsc70 and DnaJB1 without Hsp110 (**p < 0.01). Related to Figure 1.

**Figure S3.**
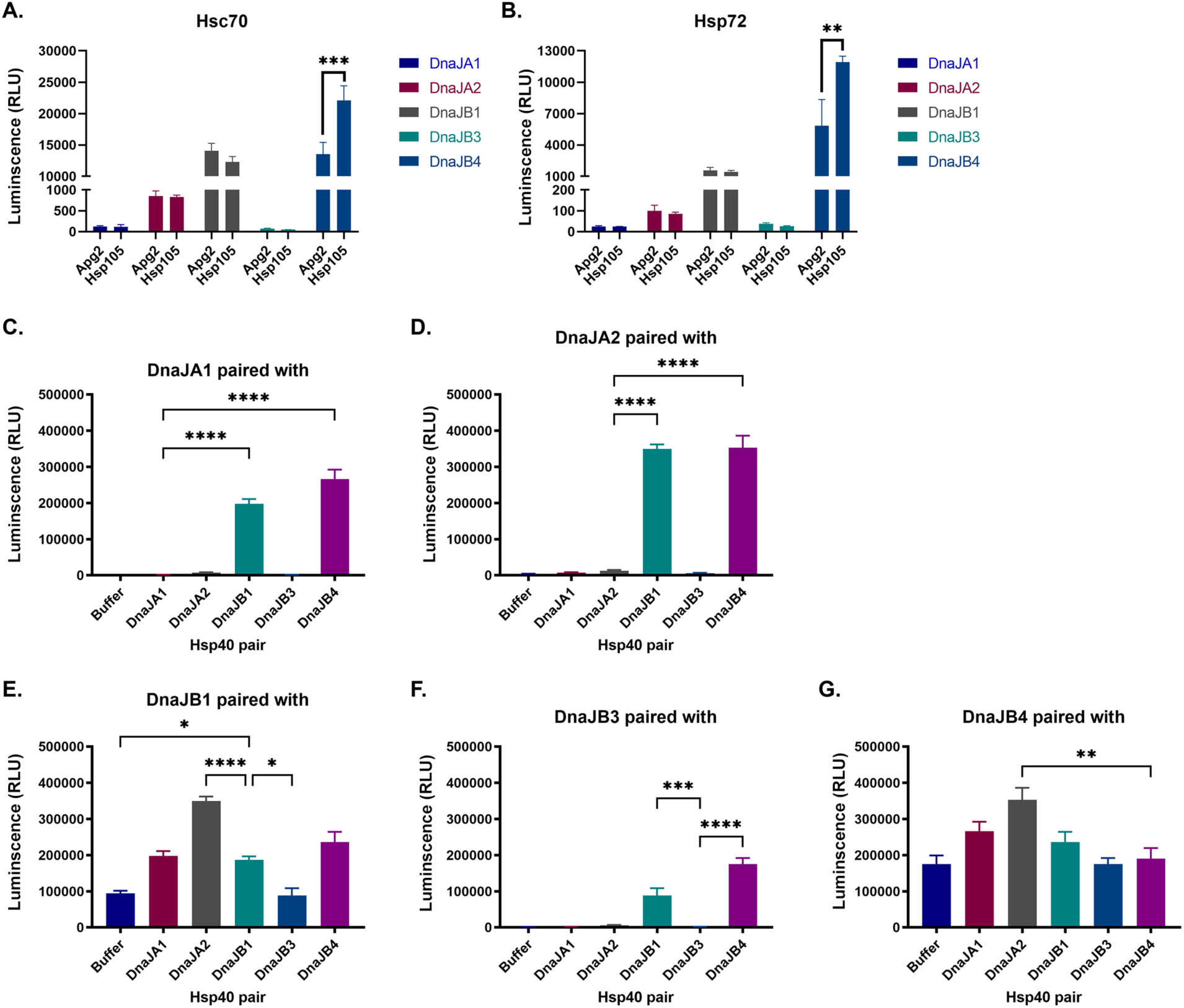
Distinct combinations of human Hsp70, Hsp40, and Hsp110 display diverse levels of protein-disaggregase activity. **(A, B)** Bar graphs showing the luciferase disaggregase and reactivation activity of every three-component combination of the (A) Hsc70 and (B) Hsp72 with each Hsp40 and Hsp110 proteins purified. 0.4μM Hsp70, 0.2μM Hsp40, and 0.04μM Hsp110 were combined with 100nM luciferase aggregates (monomeric concentration), 1% DMSO, and an ATP-regenerating system. Values represent mean luminescence ± SEM (n=4). Data were analyzed using one-way ANOVA followed by Tukey’s MCT comparing the Apg2 vs Hsp105 condition of each chaperone combination (**p < 0.01, ***p < 0.001). Data are the same as heat maps in Figure 1A and 1B. **(C-G)** Bar graphs showing the luciferase disaggregase activity of pairwise combinations of (C) 0.25µM DnaJA1 (D) 0.25µM DnaJA2, (E) 0.25µM DnaJB1, (F) 0.25µM DnaJB3, and (G) 0.25µM DnaJB4 plus either buffer, 0.25µM DnaJA1, 0.25µM DnaJA2, 0.25µM DnaJB1, 0.25µM DnaJB3, or 0.25µM DnaJB4 with Hsc70 and Apg2. 1μM Hsc70, 0.5μM Hsp40 (total, except for the buffer control, which has 0.25µM Hsp40), and 0.1μM Hsp110 were combined with 100nM luciferase aggregates (monomeric concentration) and an ATP-regenerating system. Values represent means ± SEM (n=3-10). Data were analyzed using one-way ANOVA followed by Dunnett’s MCT compared to ((A) DnaJA1 (B) DnaJA2, (C) DnaJB1, (D) DnaJB3, and (E) DnaJB4 (*p < 0.05, **p < 0.01, ***p,0.005, ****p < 0.001). Data are the same as heat map in Figure 1C. Related to Figure 1.

**Figure S4.**
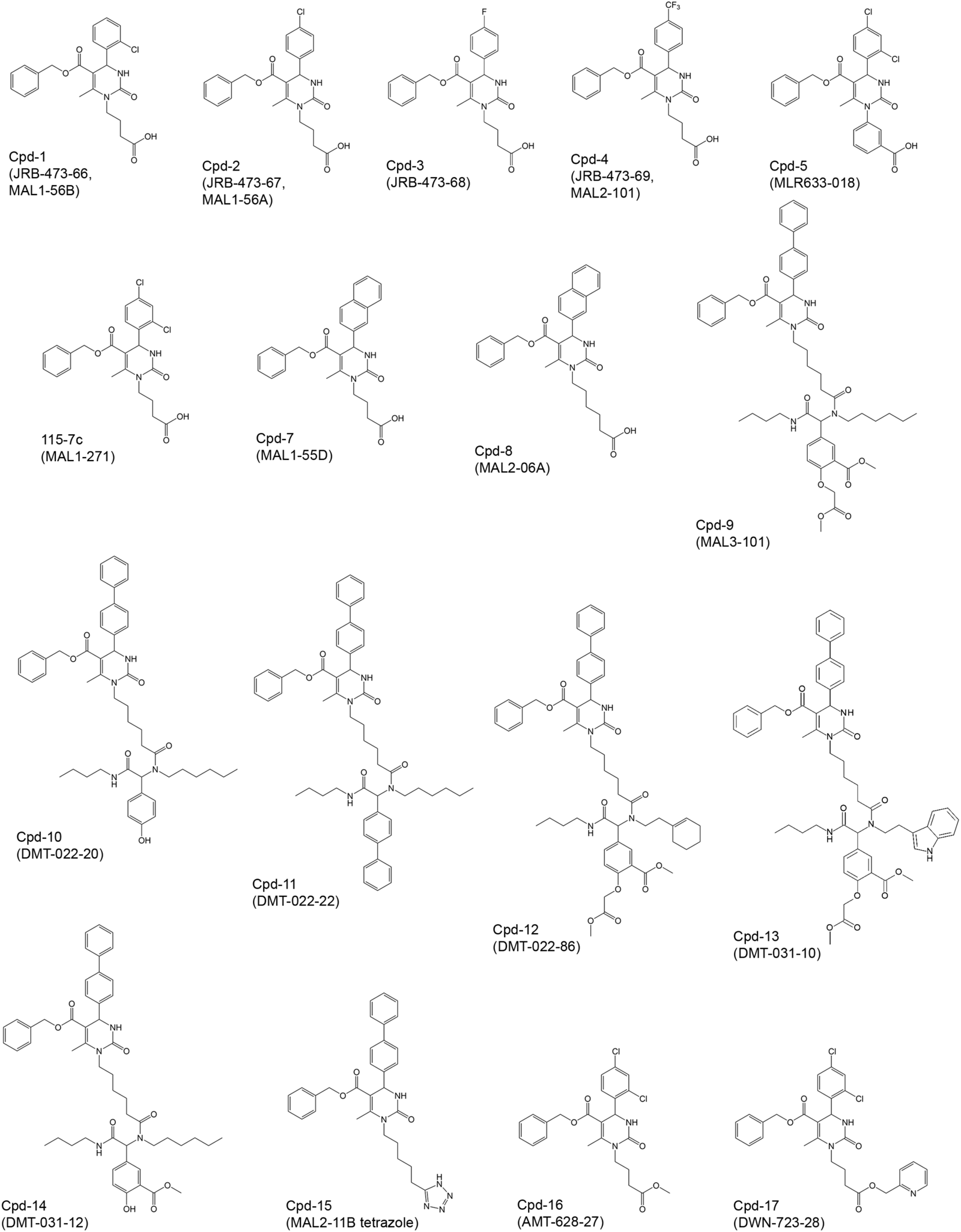
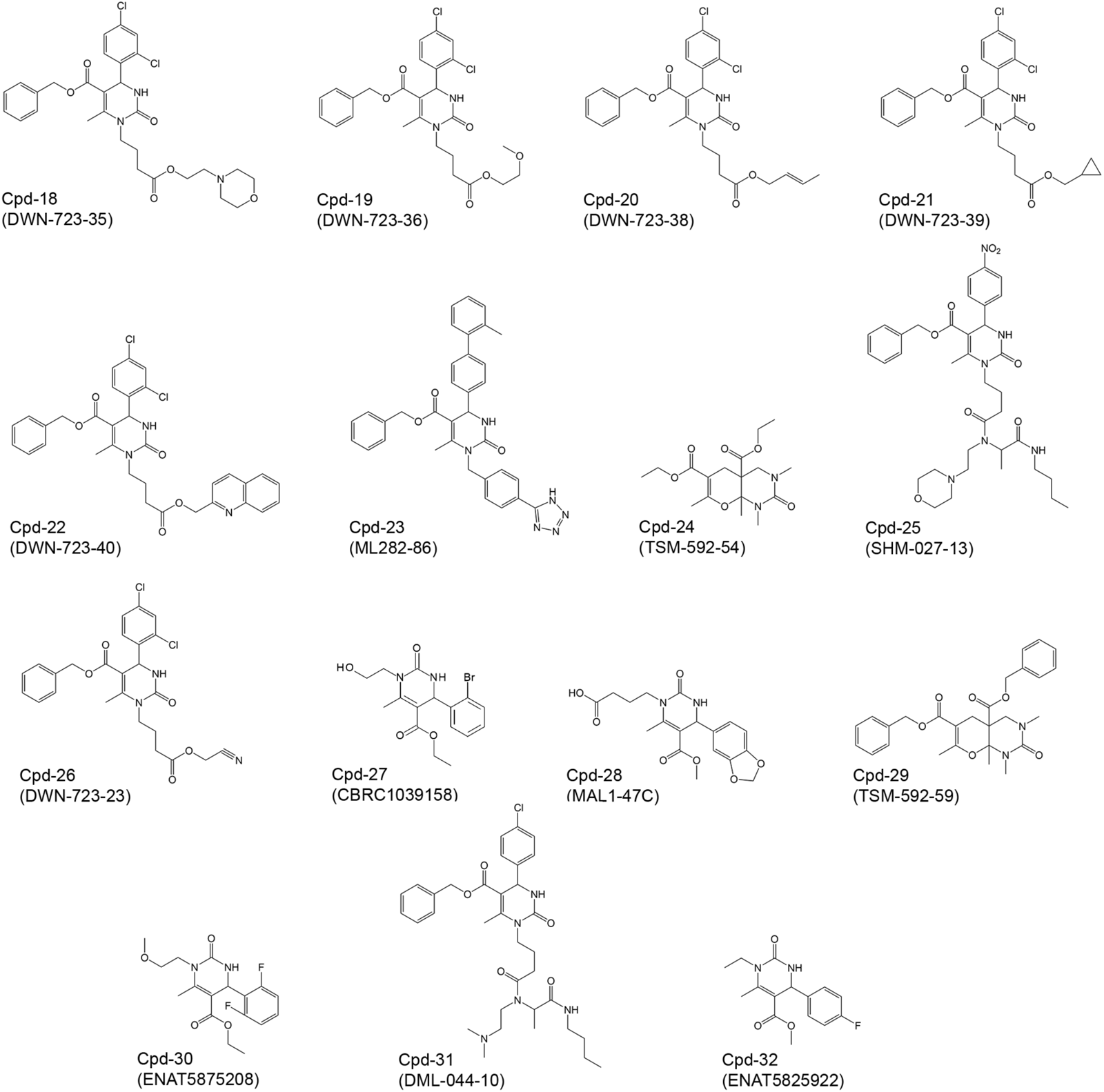
Structures of dihydropyrimidines and other compounds tested. Related to Figure 2.

**Figure S5.**
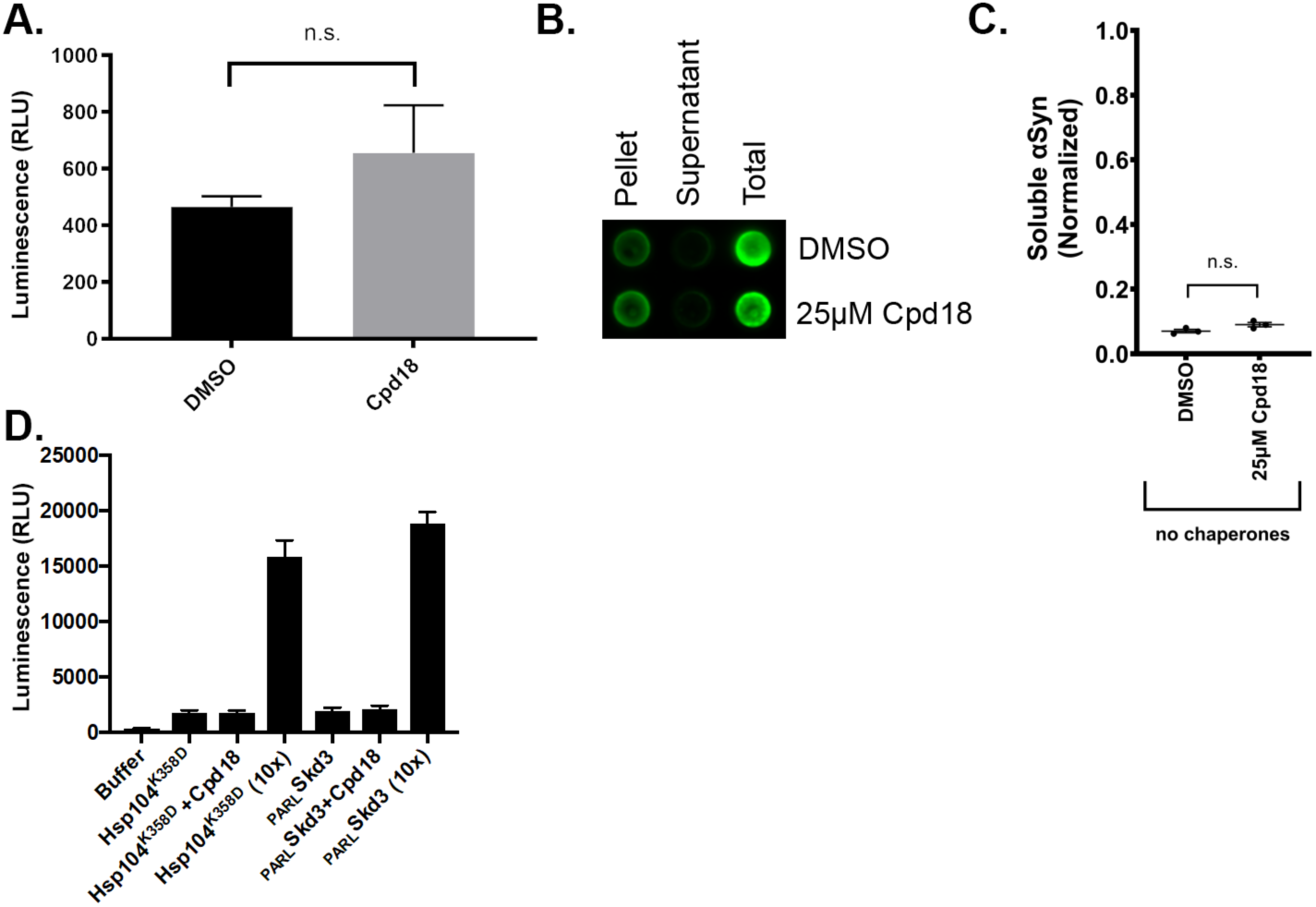
Compound 18 does not directly cause protein disaggregation or stimulate the disaggregase activity of Hsp104^K358D^ or Skd3. **(A)** 100nM luciferase aggregates (monomeric concentration) treated with an ATP-regenerating system and either DMSO or 25μM compound 18 (final 1% DMSO). Values represent mean ± SEM (n=15). Data were analyzed using unpaired t-test (ns: p > 0.05). **(B)** Representative dot blot showing αSyn content in pellet, supernatant, and total fractions after 0.5μM αSyn PFFs were treated with an ATP-regenerating system and either DMSO or 25μM compound 18 (final 1% DMSO) at 37°C while shaking at 300rpm for 90min. 10% of the total reaction, supernatant, or resuspended pellet were loaded onto the blot and stained with SYN211. **(C)** Quantification of three trials of the αSyn disaggregation assay described in (B). Dot blots were quantified using FIJI integrated density measurements. Soluble αSyn in the supernatant fraction was normalized by dividing by the total loaded αSyn for each corresponding condition and then plotted in GraphPad Prism. Y-axis represents the normalized soluble αSyn calculated. Individual data points shown as dots, bars represent mean ± SEM (n=3). Data were analyzed using unpaired t-test (ns: p > 0.05). **(D)** Luciferase aggregates (100nM monomer) were treated with buffer, Hsp104^K358D^ (0.3µM hexamer), Hsp104^K358D^ (0.3µM hexamer) plus compound 18 (10µM), Hsp104^K358D^ (3µM hexamer), _PARL_Skd3 (0.1µM monomer), _PARL_Skd3 (0.1µM monomer) plus compound 18 (10µM), or _PARL_Skd3 (1µM monomer). 1% DMSO final concentration in all samples. Luciferase disaggregation and reactivation were assessed by luminescence. Values represent mean±SEM (n=3). Related to Figure 2 and **3**.

